# Rapid evolution of fine-scale recombination during domestication: a perspective from population genomics

**DOI:** 10.1101/2024.06.05.597134

**Authors:** Zheng-Xi Liu, Ming Li, Xue-Hai Ge, Kun Wang, Si Si, Chang-Rong Ge, Jian-Hai Chen, Li-Rong Hu, Min-Sheng Peng, Ting-Ting Yin, Ali Esmailizadeh, Chang Zhang, Lu-Jiang Qu, Xue-Mei Lu, Jian-Lin Han, Ya-Ping Zhang, Ming-Shan Wang

## Abstract

Recombination serves as a pivotal force propelling the evolution of genomic diversity in sexually reproducing organisms. Despite its fundamental role, the evolutionary dynamics of recombination rates and their implications for genome evolution remain largely elusive. The domestication of animals, which is characterized by dynamic selection pressures, offers a unique lens to explore this issue. Here, we constructed a fine-scale recombination map leveraging whole-genome data from domestic chickens, encompassing both contemporary commercial breeds and the wild progenitor, the Red Junglefowl (RJF). Our analysis revealed the rapid evolution of the recombination landscape within and across chicken populations, with significant variation in recombination coldspots and hotspots, particularly in commercial breeds. The occurrence of coldspots and the disappearance of hotspots are closely associated with selective sweeps.Contrary to prevailing observations in most species, we observed a weakly negative correlation between recombination rates and the frequency of introgressed ancestry from other RJF subspecies. Our findings provide valuable insights into the intricate interplay of evolutionary forces shaping the evolution of recombination.

## INTRODUCTION

Meiotic recombination is a crucial process in which DNA exchange occurs between homologous chromosomes, ensuring their accurate segregation in sexually reproducing species. This process is central to generating and maintaining genetic diversity, facilitating the development of novel haploid gametes. Empirical evidence reveals that recombination serves as a major force in both micro- and macroevolutionary processes (Palahi et al. 2023; Paigen and Petkov 2010), driving the formation of new combinations of alleles and haplotypes. This enables the efficiency of adaptive evolution and selection, highlighting its practical and fundamental significance. Investigating the recombination landscape provides insights into how recombination influences the selection, evolution, and demographic history of populations. Moreover, it allows for a broader exploration of the mechanisms and forces shaping the evolution of genomes and species (Fransz et al. 2016; Marand et al. 2019; Dutta et al. 2017). Practically, manipulating the recombination rate has practical implications, potentially enhancing the efficiency of the breeding processes (Battagin et al. 2016; Epstein et al. 2023).

Influenced by demographic, environmental, and genetic factors (Stapley et al. 2017; Dreissig, Mascher, and Heckmann 2019; Hoge et al. 2024; Jones et al. 2019), the recombination rate is not static but exhibits high variation across taxa, populations, and individuals, even within chromosomes(Stapley et al. 2017; Penalba and Wolf 2020). This rate is unevenly distributed across the genome in many species, with higher rates found predominantly located at the distal ends of chromosomes, termed “hotspots”, and lower rates (coldspots) found in the pericentromere (Paigen and Petkov 2010). These hotspots are typically 1-2kb in size (Paigen and Petkov 2010). In mice and humans, the position of the hotspot is largely determined by *PRDM9*, a zinc-finger protein that binds to specific DNA motifs (Hochwagen and Marais 2010). However, the binding domain of *PRDM9* evolves rapidly, and hotspot regions show a rapid turnover (Auton et al. 2012; Brick et al. 2012; Hinch et al. 2011; Hoge et al. 2024). Despite lacking functional *PRDM9*, species like birds, canids and yeast maintain recombination patterns and hotspots (Singhal et al. 2015a; Axelsson et al. 2012; Lam and Keeney 2015). The locations of recombination hotspots tend to occur at promoters or near transcription start sites (TSSs), remaining highly stable over evolutionary time (Singhal et al. 2015b). However, the molecular basis influencing the distribution and occurrence of recombination events is poorly understood in many organisms, particularly those lacking functional *PRDM9*.

Recombination can influence the efficacy of selection, and reciprocally, it can be influenced by episodes of selection (Roze 2021). Domestication is a long-term experiment by humans with animals and plants over thousands of years, involving intense selection and breeding for targeted traits that have shaped the genetic architecture and physiological and phenotypic diversity of domesticated species. This provides a unique opportunity to investigate the interplay between the selection and recombination. However, existing studies have yielded inconsistent and sometimes contradictory results. Some suggested an overall increase in recombination rates during domestication, positing that selection generally favors increased recombination during periods of rapid evolutionary change (Otto and Barton 1997; Ross-Ibarra 2004). While others argue that higher recombination is a preadaptation to domestication. In this view, a population with a higher recombination rate would increase its efficiency and responsiveness to the selection, making it more likely to be successfully domesticated (Gornall 1983). Additionally, Lenormand and colleagues proposed that a reduced recombination rate in domesticates may confer benefits by protecting against maladaptive gene flow from wild relatives (Lenormand and Otto 2000).

The evolution of recombination rates through domestication remains a topic of debate, with most genome-wide comparisons focused on plant species rather than animals. Animal domestication episodes and processes may differ from those in plants (Larson and Burger 2013; Larson et al. 2014). Previous investigations into the relationship between recombination and domestication vary in the datasets and approaches, and the types of wild and domesticated breeds compared (Ross-Ibarra 2004; Munoz-Fuentes et al. 2015). Notably, some studies have not included the direct wild ancestors of the domesticated species for comparison. One reason is that the direct wild ancestor of many domesticated species such as horses and cattle, are extinct (Larson and Burger 2013; Larson et al. 2014). Directly comparing the recombination landscapes between these domesticated species and the wild populations from which they were not directly derived may introduce bias when assessing the impact of domestication on recombination (Ross-Ibarra 2004). As a result, how domestication and continued selection affect the evolution of recombination rates remains controversial.

Chickens have become the most abundant domesticated species. Unlike many other domesticated animals, their direct wild ancestor, the Red Junglefowl (RJF), still exists in South and Southeast Asia. Our earlier study demonstrated that chickens trace their origins to the RJF subspecies *Gallus gallus spadiceus* (GGS) and that the chicken domestication process, was marked by intense and dynamic selection and breeding, which led to a significant bottleneck and the accumulation of deleterious mutations (Wang et al. 2021). Domestic chickens exhibit lower genetic diversity and an increase in linkage disequilibrium (LD) over longer distances compared to RJF (Wang et al. 2021), suggesting alterations in recombination rates and patterns post-domestication. Additionally, as chickens spread globally, they engaged in hybridization with other jungle fowl species, acquiring additional ancestry that likely modified their genetic architecture. Domestic chickens and GGS therefore provide a valuable system with which to investigate the intricate relationship between domestication and recombination landscapes at finer scales (Wang et al. 2020; Wang et al. 2021).

In this study, we explored recombination rates and profiles by analyzing over 120 whole nuclear genomes of GGS and domestic chickens, including both village chickens and recently improved commercial breeds. We used a population genomic approach to construct the landscape of recombination for GGS and domestic chicken breeds. We then characterized the variation of recombination landscape, identified recombination hotspots and hotspots, and investigated their genomic features and motifs as well as the associations with selection, and gene flow. This study offers insights into the landscape of recombination variation in domestication and enhances our understanding of how domestication and selective breeding affect the evolution of recombination.

## RESULTS

### The fine-resolution recombination landscape of chicken

We generated whole-genome sequences from 17 GGS subspecies of RJF, 20 Jining Bairi chickens (JNBR, a local Chinese breed from Shandong province), and 20 White Leghorns (WL, a commercial egg-laying breed originating in Italy). The average sequencing depth for each genome was 16.46× (ranging from 7.89× to 26.14×). We also collected 64 genomes from previous studies and combined with our dataset, creating a comprehensive dataset of 121 genomes (supplementary Table S1). This dataset encompasses diverse populations, including GGS, four local domesticated chicken (LDC) breeds, namely Chahua (CH, a native breed from south-western China with similar phenotypes and behavior to the RJF), Nixi (NX, an indigenous Chinese chicken breed from Yunnan province that is highly adapted to living in a cold climate), JNBR, and Luxi Fighting (LX, a Chinese game fowl breed from Shandong province) chickens, as well as a modern commercial breed WL. Each population is represented by 20 samples. A green jungle fowl (*Gallus varius,* GV) genome was used as an out-group. These chicken breeds include the ancestral variety from the region that is closely related to the putative center of chicken domestication, as well as the two clades of modern chickens from a panel of global chickens identified in our previous diversity study (Wang et al. 2020).

Following quality control filtering, we identified 7,203,349 (WL) to 14,171,854 (GGS) SNPs per population, corresponding to 7.63 to 15.00 SNPs per kilobase (kb) (supplementary Table S2). The maximum likelihood tree (supplementary Fig. S1) illustrated that domestic chickens and GGS clustered distinctly into separate groups, with GGS positioned basally to LDC. An exception is WL, which occupies a basal position relative to GGS and LDC, which is likely due to admixture and carrying high levels of additional ancestry from another divergent RJF lineage, the *G. g. murghi* (GGM) (Wang et al. 2020). Compared to GGS, LDC and WL exhibited lower genetic diversity, and greater LD decay distance (supplementary Fig. S2, S3), which is consistent with prior findings that chickens have undergone selective bottlenecks since domestication (Wang et al. 2021).

We used RelERNN, a deep learning method, to construct a high-resolution relative recombination rate map from polymorphism data (Adrion, Galloway, and Kern 2020). Both the number of training runs and the sample size could affect the results (Adrion, Galloway, and Kern 2020). To assess the robustness of the results across different parameters, we first tested the effect of the number of training runs by performing 100, 200, 500, 1000, and 2000 simulated training runs under an equilibrium demographic scenario. With more than 500 training runs, the correlation coefficient of the predictions exceeded 0.90 (supplementary Fig. S4). Despite additional training (e.g., 1000 and 2000), the correlation only marginally increased, suggesting proximity to saturation. Considering both the consumption of computational resources and the accuracy of the results, we conducted 1000 training sessions in subsequent analysis. Then, to explore the effect of sample size on the estimation, we estimated the recombination rate under an equilibrium demographic scenario using 5, 10, and 20 samples and found that the estimates showed a high degree of consistency when the sample size was equal to or greater than 10 (Pearson’s r = 0.93; *P*-value < 2.2e-16, supplementary Fig. S5). Hence, 20 samples per population are sufficient to reconstruct their recombination landscapes. Taken together, these analyses and tests provide a solid justification and foundation for the use of RelERNN to estimate the recombination map.

Considering the differences in genetic background and demography between populations, we used SMC++ (Terhorst, Kamm, and Song 2017) to infer the demographic history of each population. The effective population size (Ne) varied between populations. GGS has the largest Ne in the recent past, while WL had the smallest Ne (supplementary Fig. S6). Then, we estimated the recombination rate for GGS and each of the five domestic chicken populations using the equilibrium scenario and the SMC++ inferred demographic scenario, respectively. The magnitude of the inferred recombination rates under the two scenarios is consistent, and the Pearson’s correlation between these estimates ranges from 0.27 (LX) to 0.78 (GGS) (supplementary Fig. S7). Given the moderate correlation between the estimates under the two scenarios, our subsequent analyses were based on the estimates under the SMC++ inferred demographic scenario.

We revealed remarkable variations across populations and chromosomes. Among the six populations, the genome-wide average recombination rate was estimated to be 1.13e-09 c/bp (crossovers per base pair), ranging from 9.01e-10 c/bp (WL) to 1.27e-09 c/bp (GGS). Our estimate is consistent with previous estimates for focal passerine birds (Groenen et al. 2009; Provost et al. 2022), suggesting the robustness of our analysis.

### The distribution and pattern of the recombination rate

One of the striking characteristics of the avian genome is the distinguished length differences between chromosomes, which is also reflected in many genomic features, e.g. GC content, genetic diversity, gene density, as well as recombination rate (Groenen et al. 2009; Fullerton, Bernardo Carvalho, and Clark 2001). We observed a trend emerging with microchromosomes displaying a higher recombination rate than macrochromosomes (mean 1.18e-09 c/bp for microchromosomes and 1.11e-09 c/bp for macrochromosomes). The average Pearson’s correlation coefficient for recombination rate (c/bp) versus chromosome length (in bp) ranges from −0.25 to −0.56 (supplementary Fig. S8). We compared the level of recombination with respect to several genomic features, including GC content, genetic diversity, and gene density, and found a positive association between recombination rate and these features (supplementary Fig. S9). Moreover, compared to introns and intergenic regions, exons exhibit a higher recombination rate, with the promoter regions displaying elevated rates compared to other regions (supplementary Fig. S10). This result is in line with previous studies in the zebra finch and long-tailed finch (Singhal et al. 2015a), monkeyflower (Hellsten et al. 2013) and tomato (de Haas et al. 2017), and supports the notion that recombination has been primarily localized to the promoter regions of genes in birds and has also remained highly conserved over millions of years (Singhal et al. 2015a).

Prior studies propose a “U”-shaped distribution of recombination rates across the chromosomes in mammalian species, such as mice and humans, with recombination rate tending to be increased near telomeric regions and suppressed in centromeric regions (Stevison et al. 2016; Spence and Song 2019; Nowosad et al. 2020). However, in chickens, we did not find an apparent increase in the recombination rate towards distal positions for each autosome. Instead, the distribution of recombination rate exhibited high variation between chromosomes (Fig. 1A). For example, the recombination rate across chromosome 1 displayed a relatively uniform distribution, without a clear central region of decrease and an increase in the distal regions. This result is slightly inconsistent with the previous study (Groenen et al. 2009), but aligns more closely with a recent study (Weng et al. 2019), likely due to the later study employing a larger set of genetic markers and a more complete reference genome. The improvement in the reference genome is likely to be due to the improved assembly of microchromosomes and subtelomeric regions, which may improve the efficiency and accuracy of genotyping in these regions. Meanwhile, some chromosomes (e.g., chr6, chr8) showed a central region of reduced recombination and an increase in the distal regions, akin to observations in humans and mice (Wooldridge and Dumont 2023; McVean et al. 2004; Spence and Song 2019).

**Fig. 1.**
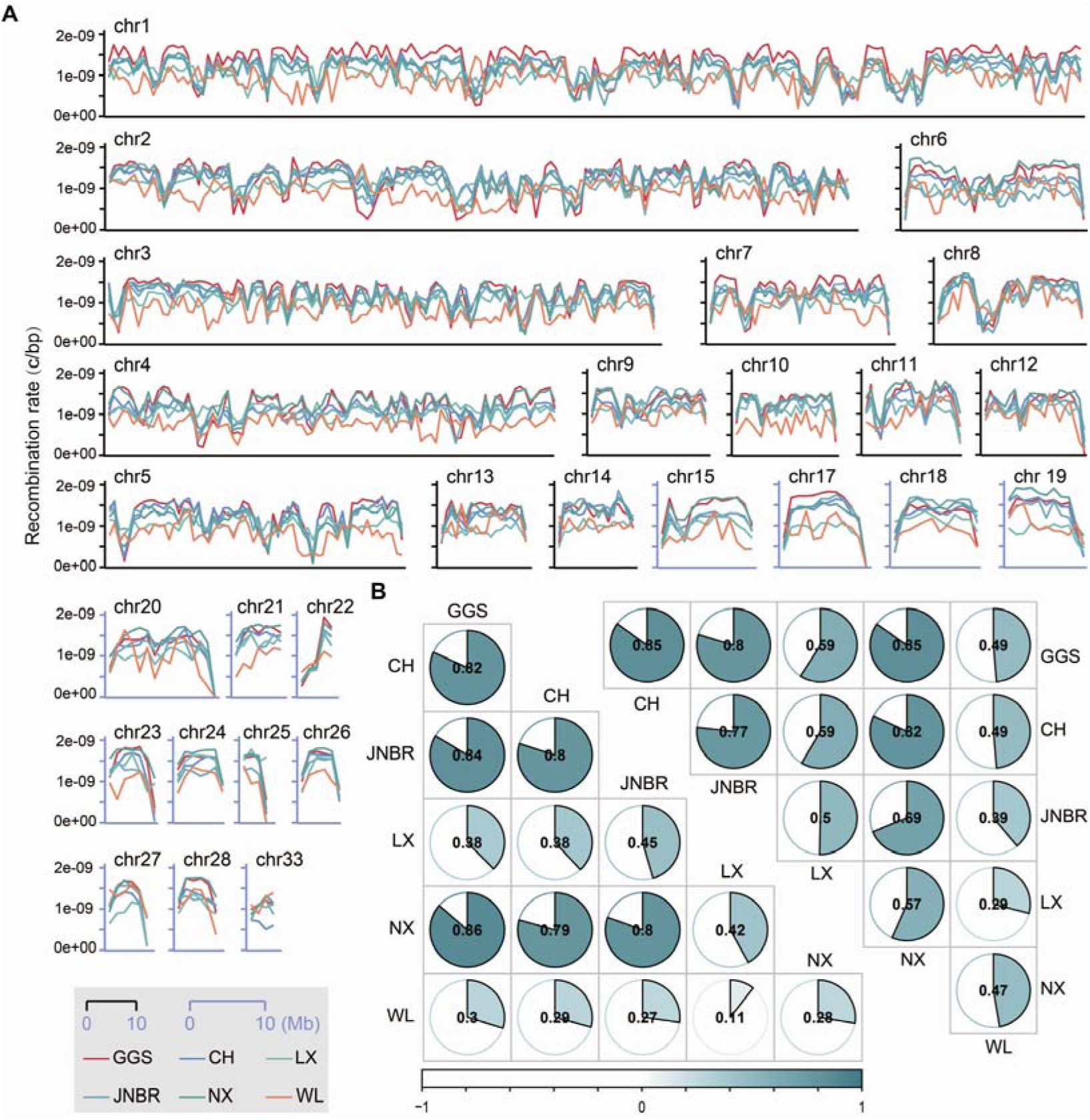
Landscape of recombination rates in GGS and domestic chickens. (A) The distribution of the recombination rate (c/bp). Estimates are normalized using 1-Mb sliding windows. (B) Pearson’s rank correlation coefficient for the recombination rate for macrochromosomes (left) and microchromosomes (right) from each pair of populations. Macrochromosomes include chr1-5 and the rest are microchromosomes. Dot chromosomes were excluded from this analysis due to their limited length and high missing rate in variant calling. GGS, *Gallus gallus spadiceus*; CH, Chahua chicken; JNBR, Jining Bairi Chickens; LX, Luxi Fighting; NX, Nixi; WL, White Leghorn.

On a broader scale, macrochromosomes and microchromosomes exhibited high differences in recombination landscapes. Microchromosomes displayed significant variability in the distribution of the recombination rate. Some (e.g., chr17, chr18, chr23) exhibited unimodal distribution of the recombination rate, with peaking in the central regions and a progressive decline towards the ends. On others (e.g., chr21, chr22), recombination decreased at one end but progressively increased towards the opposite end. Additionally, certain chromosomes exhibited a central region of reduced recombination, alongside reductions in subtelomeric regions. This is a prevalent (though not exclusive) pattern in medium-sized chromosomes (e.g., chr15, chr20). However, this may not necessarily indicate a distinct recombination pattern at the telomeres of each chicken chromosome, due to incomplete assembly (Li, Sun, et al. 2022). Nevertheless, a similar pattern is evident in the previous version of the chicken recombination map (Groenen et al. 2009) and in other species such as butterflies (Palahi et al. 2023).

### Rapid evolution of recombination rate in different populations

To compare the broad-scale recombination map between different populations, we first binned the genomes with 1Mb windows and calculated the Pearson’**s** correlation coefficient (r) for the recombination rate from each pair of the populations (Fig. 1B, supplementary Fig. S11). The Pearson’**s** correlation coefficient ranges from 0.20 to 0.84, suggesting a rapid evolution of recombination on the broad scale. The phylogenetic relationships between these populations cannot be fully reconstructed from the recombination rate (supplementary Fig. S12), which suggests that recombination rates evolve dynamically between different populations.

Microchromosomes display a higher correlation than macrochromosomes (Fig. 1B), aligning with greater conservation of microchromosomes over macrochromosomes in avian species (Waters et al. 2021). The populations within LDC show a higher pairwise correlation (Pearson’s r ≥ 0.46) than the WL. Of particular note, WL displays the lowest correlation coefficient of recombination rate compared to domestic chicken breeds and GGS (supplementary Fig. S11), aligning with the result that WL has the lowest genetic diversity and largest LD decay over distance (supplementary Fig. S2, Fig. S3). These findings underscore the significant impact of recent breeding and selective bottlenecks in shaping the genetic architecture landscape of WL.

The domestic population exhibits a lower recombination rate than the GGS, which exhibits the highest rate (Fig. S13). To test whether this pattern extends to other domestic species, we reanalysed the genomes of ducks, goats, pigs and sheep, alongside those of their wild counterparts. Three domestic populations were included per species (see Supplementary Table S3). In all cases, the domestic breeds exhibited lower recombination rates than their wild relatives (see Fig. S13), which is consistent with the trend observed in chickens. However, since domestication and selective breeding result in a loss of genetic diversity and an increase in homozygosity, estimates of recombination may be biased downward. While some studies show a decline in the recombination rate in maize (Hufford et al. 2012), tomatoes (Fuentes et al. 2022) and some domestic animals (Larson and Burger 2013), our study can not to clarify whether domestication has resulted in a reduction in recombination compared to wild populations.

### Characterize recombination hotspots and coldspots

Recombination hotspots exhibit a diverse distribution in eukaryotic organisms that have *PRDM9* (Singhal et al. 2015a). To investigate how selection influences the evolution of recombination hotspots, we used a previously described method (Palahi et al. 2023) to identify recombination hotspots in each chicken breed using a 1 kb window size by comparing with the background. We identified a total of 128,305 hotspots across all six breeds, with an average of 21,384 per population (Fig. 2A-B), which are distributed non-uniformly across the chicken genome. Although differing in approaches, the number of hotspots in chickens is comparable to estimates for mammals. For example, 25,000-50,000 have been reported in human populations (Myers et al. 2005).

**Fig. 2.**
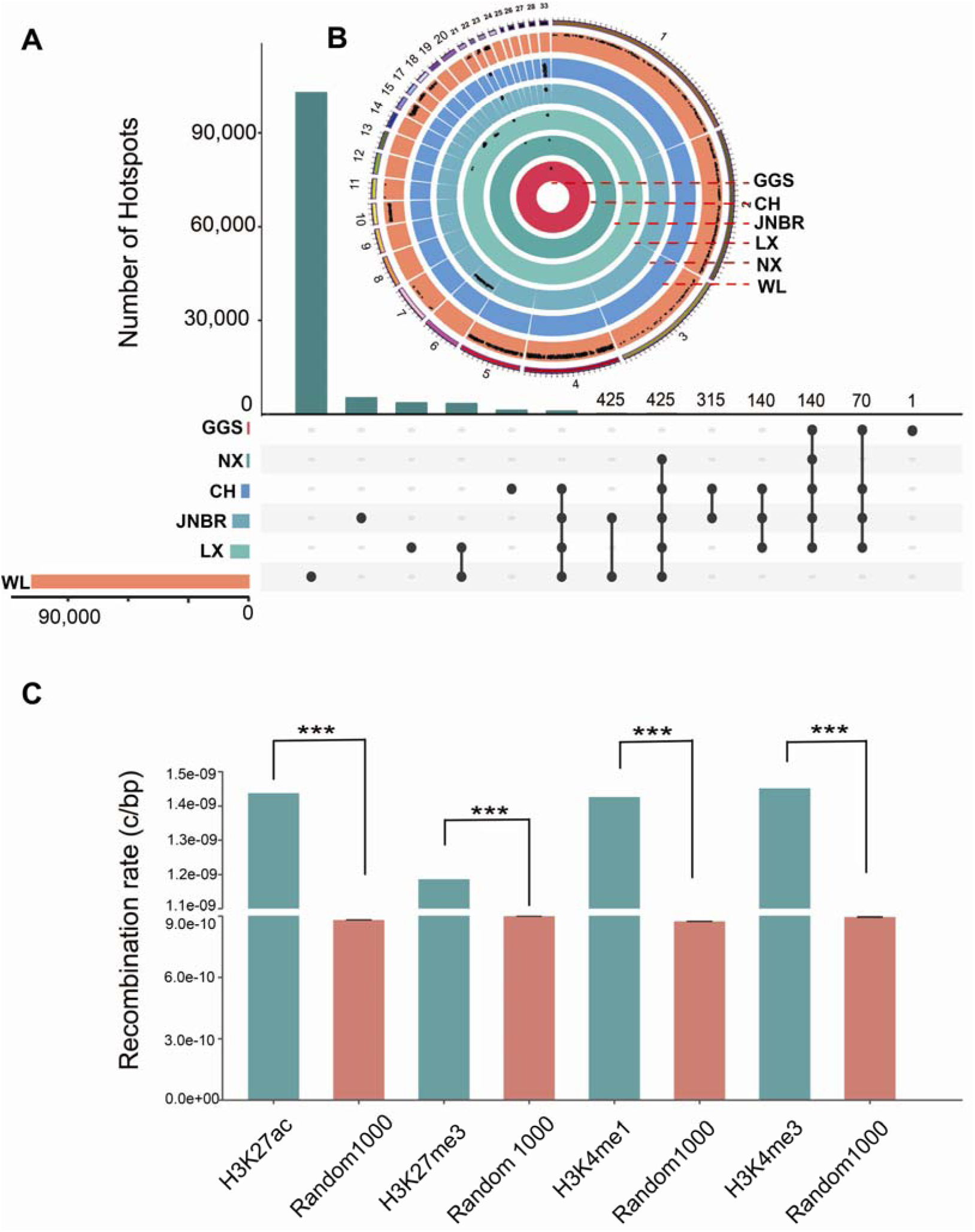
Recombination hotspots variation across genomes. (A) Identification of shared and breed-specific recombination hotspots in GGS and chickens. (B) The localization of recombination hotspots on each chromosome is depicted in concentric circles, with GGS, NX, LX, JNBR, CH, and WL represented from innermost to outermost circles. (C) Comparison of recombination rates between regions associated with four histone modifications (H3K27ac, H3K27me3, H3K4me1, and H3K4me3) and randomly selected regions (Random_1000, n=1000) of equal length from the rest of the genome. *** indicates P < 0.001 by one-sample t-test.

Prior examination of mammals, including humans and mice, demonstrated a considerable influence of the functional genomic elements on local recombination rates (Brick et al. 2012; Spence and Song 2019). In our study, we reanalyzed ChIP-seq data for 4 histone marks including H3K4me3, H3K4me1, H3K27ac, and H3K27me3 reported in our previous study (Pan et al. 2023) and measured the recombination rate for these histone-marked regions. Our findings reveal variations in recombination rates among different chromatin states. Regions associated with H3K4me3, H3K4me1, H3K27me3 and H3K27ac showed significantly higher recombination rates than expected by randomly selected regions in each chicken population (*P*-value < 0.001, n=1000 times; Fig. 2C, Fig. S14), with the exception of the H3K27me3 in WL. Our result suggests that the recombination event usually accompanies the histone chromatin modifications, consistent with prior research indicating that double-strand breaks (DSBs) tend to occur in regions proximal to histone modifications in *PRDM9* knockout mice (Brick et al. 2012). Although a causal relationship has yet to be established, this suggests that these histone modifications are also necessary to create a favorable chromatin environment for the initiation of meiotic recombination in birds. It is noteworthy that both recombination rates and chromatin structure are susceptible to temporal changes. Our recombination estimates offer a historical average for each population, whereas chromatin state modifications are derived from contemporary samples. This inconsistency may potentially introduce limitations to the analysis, despite the prior comparison between them (Spence and Song 2019).

### Recombination hotspots and coldspots evolve rapidly

We compared the number of shared hotspots among chicken populations and observed remarkable variation (Fig. 2A). Thousands of hotspots were unique to specific populations, with only 425 hotspots shared by all domestic populations. Domestic populations shared more hotspots with other domestic populations than with GGS. Furthermore, we examined the frequency of recombination hotspots and coldspots overlapping with selective sweeps. The overlap between the recombination hotspots and selective sweeps was approximately 2.25%, which is less than expected by chance when compared to the randomly selected regions (approximately 11.32 ± 0.33%, one-sample t-test, *P*-value < 0.001), suggesting that most of the selective sweeps occur outside the hotspot. In contrast, the overlap between coldspots and the selective sweeps (approximately 24.49%) exceeds the expected value compared to the randomly selected regions (approximately 2.37 ± 0.16%; one-sample t-test, *P*-value < 0.001).

It is important to note that although empirical approaches are commonly used to find selective sweeps, they may introduce bias and cannot rule out false positives. To further explore our analysis, we systematically retrieved putatively selective sweeps with progressively more relaxed empirical cut-offs (from the top 1% to 5% and 10%), expecting an increase in false positives. Consistent with our expectation, the proportion of overlap between coldspot and putatively selective sweeps progressively decreases (from 24.49% to 15.75% and 10.31%), and the frequency between hotspot and selective sweeps increases progressively (from 2.25% to 7.59% and 9.04%; supplementary Fig. S15). This suggests that selective sweeps tend to occur in or favor regions of low recombination.

### Identify the recombination hotspot determinants by deep learning method

In humans, the DNA-binding protein PRDM9 is the factor that controls the localization of recombination hotspots by specifically binding to the 13-bp consensus DNA motif: CCNCCNTNNCCNC (Myers et al. 2005). To examine whether chickens lacking functional *PRDM9,* possess DNA-binding motifs and sequences associated with hotspots, we leveraged RHSNet (Li, Chen, et al. 2022) to predict the potential determinant motifs of recombination hotspots in each chicken population (Fig. 3A). Across all six chicken populations, the GC content of the top 100 predicted motifs is significantly higher than that of the entire genome (56.93% vs. 45.43%; Fig. 3C). This is in line with previous studies that the recombination hotspot often occurs at CpG islands (CGIs), and genomic regions with elevated GC content.

**Fig. 3.**
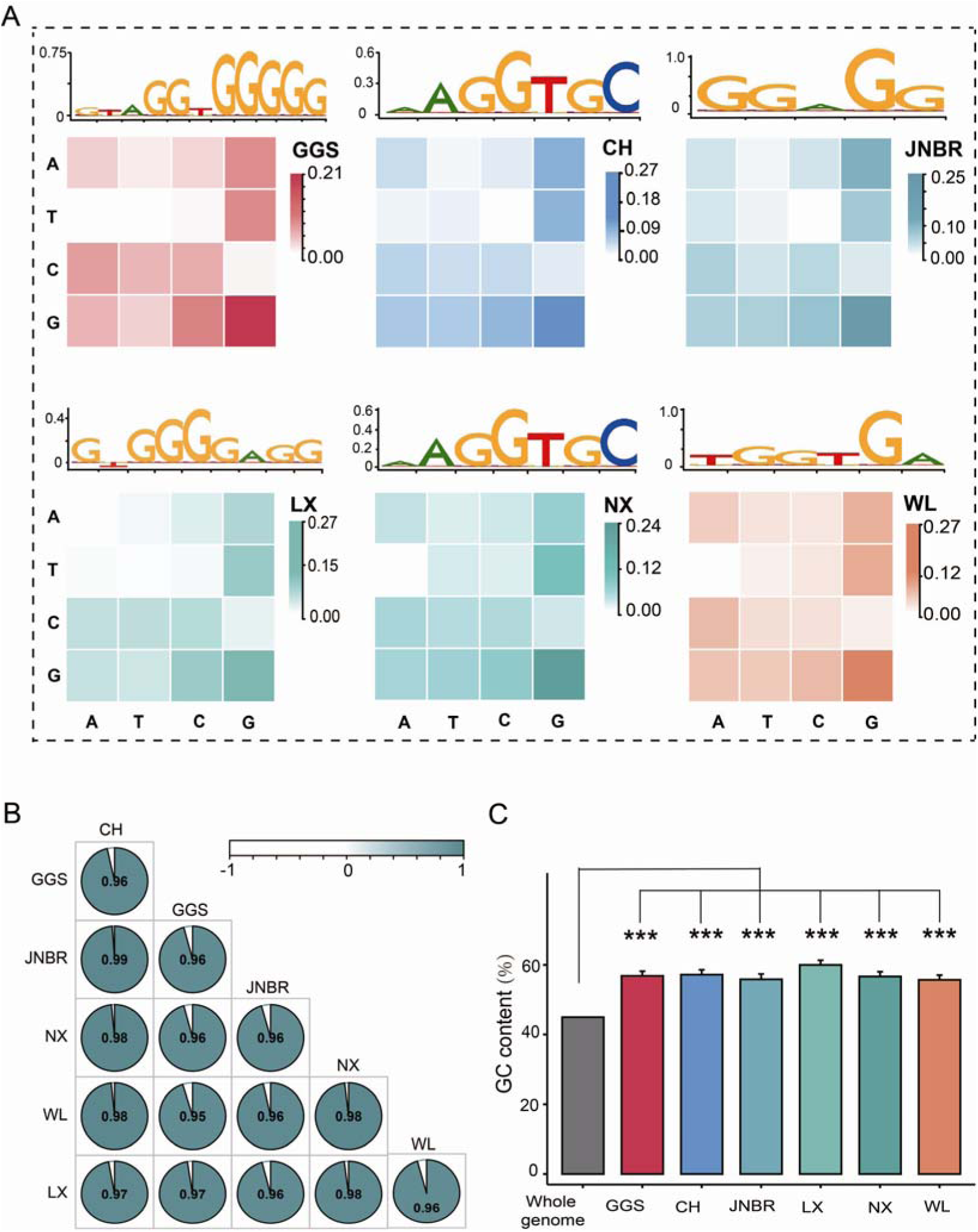
The distribution of the predicted motif for the recombination hotspot across different populations. (A) Results of the composite motif detected by RHSNet. The top shows the motif embeddings in each chicken population, and the bottom heatmap indicates the frequency of 2-mer occurrence in the detected motifs. (B) Pearson’s correlation coefficients between the frequency of 2-mer motif between each pair of populations. (C) Average GC content in the whole genome and motif sequences. *** indicates P < 0.001 by one-sample t-test.

To provide a detailed representation of the motif embeddings, we followed a previous study (Li, Chen, et al. 2022) to count and visualize the motifs in 2-mer occurrence frequency (Fig. 3A). The occurrence frequency of the 2-mer motif exhibits a high correlation among each population (Pearson’s r ≥ 0.95; Fig. 3B). The 2-mer motifs in WL exhibited a stronger correlation with other populations (average Pearson’s r = 0.97) than the correlation observed in broad-scale recombination between WL and the other population (average Pearson’s r = 0.33). Notably, the {GG}_n_ motif exhibited extremely high enrichment and had a high frequency in each population. This finding suggests that, while the location of the hotspot varies considerably, the binding motif for the recombination determinant remains relatively conserved in recently diverged populations.

### The correlation between recombination rate and introgressed ancestry

Hybridization with closely related wild species commonly occurs in domesticated species as they expand from their origin center (Quilodran, Montoya-Burgos, and Currat 2020). These introgressed materials often facilitate the adaptive evolution of domesticated species (Ma et al. 2019; Zheng et al. 2020). The recombination rate plays a crucial role in determining the pattern and frequency of introgressed ancestry (Kim, Huber, and Lohmueller 2018). Positive correlations between the recombination rate and the level of introgressed ancestry have been observed in many organisms, such as swordtail fish (Schumer et al. 2018), house mice (Janousek et al. 2015) and butterflies (Martin et al. 2019), which have been assumed to purge of deleterious mutations within introgressed regions because of incompleteness. To investigate the relationship between the amount of introgression and recombination rate in chickens, we focus on WL, a commercial breed that hybridized with GGM (diverged from chickens ∼80 kya) and then inherited over 10% of its genetic ancestry from GGM (Wang et al. 2020). Strikingly, the frequency of introgressed ancestry showed a weak negative correlation with recombination rate (Fig. 4, supplementary Fig. S16), regardless of whether Loter (Dias-Alves, Mairal, and Blum 2018) or fd (Martin et al. 2019) was used to identify GGM introgressed ancestry.

**Fig. 4.**
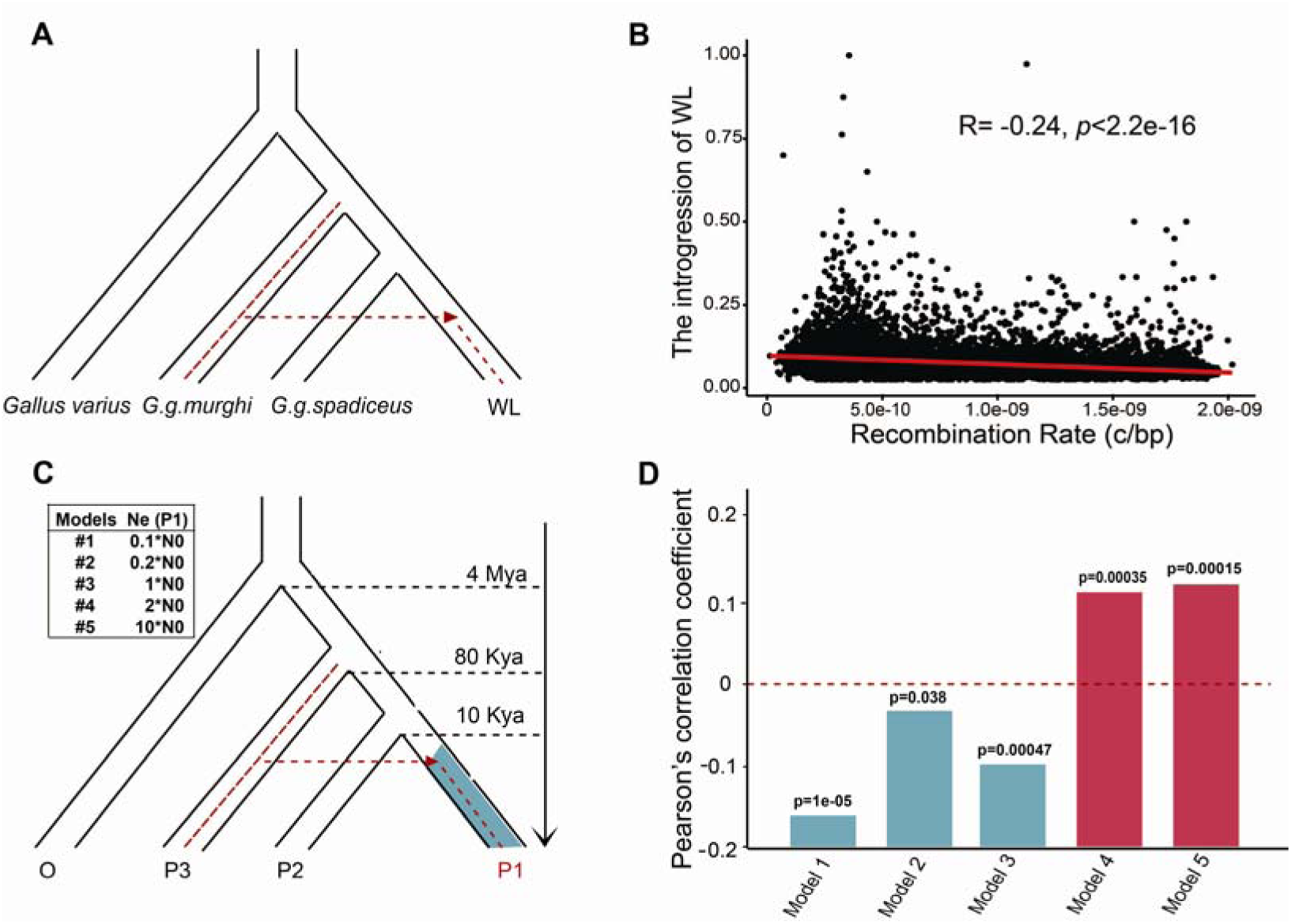
Correlation between introgression frequency and recombination rate. (A) The population topology for the detection of GGM introgressed regions in WL. (B) Correlation between the recombination rate and the frequency of introgression. (C) The topology for the four populations used in the simulations. In each of the five models, the population size for O, P3, and P2 is set to N0, and the population size for P1 is set to 0.1*N0, 0.2*N0, N0, 2*N0, and 10*N0, respectively. (D) Pearson’s correlation coefficients between the frequency of gene flow and the recombination rate under five models.

We further examined the overlap proportions between introgressed regions and recombination coldspots/hotspots. The overlap between coldspots and introgressed regions, is significantly higher than expected by chance from randomly selected regions (16.27% vs. 2.38%; permutation test, *P*-value < 0.001). In contrast, the overlap between recombination hotspots and introgressed regions is significantly lower than expected by chance from randomly selected regions (9.89% vs. 11.29%; permutation test, *P-*value < 0.001). Interestingly, when a statistically stricter cut-off was used to retrieve the introgressed regions, a higher proportion of overlap was found between coldspots and these introgressed regions (supplementary Fig. S17). This result aligns with our above finding that introgressed ancestry in WL is more likely to be obscured in regions of low recombination, contrary to empirical investigations of hybridization and recombination rates in the majority of species.

Demography, such as population size fluctuations and introgression, could influence recombination patterns and genetic dynamics (Simons and Sella 2016; Ritz, Noor, and Singh 2017; Kim, Huber, and Lohmueller 2018). As a commercial breed, WL has experienced continuous selection bottlenecks and maintained a long-term small population size (Wang et al. 2021). We speculate that the long-term small population size of WL may contribute to the unexpected negative correlation. To test this, we performed a simulation analysis. Our models considered the demographic history of chickens and RJF (Wang et al. 2020; Wang et al. 2021) and simulated the scenarios with four populations having a topology (((P1, P2), P3), O) reflecting the phylogeny and splitting of GV, GGM, GGS, and WL (Fig. 4C) where the P3 (donor), with a splitting of 80 thousand years from P1 and P2, contributed 10% of the gene flow to the recipient population (P1). We assumed population sizes for the recipient population with different sizes and measured the overlapping frequency of gene flow and recombination rate. When the effective population size for the recipient population was larger (e.g., 2 times and 10 times) than the donor population, the frequency of gene flow was positively correlated with the level of recombination rates in scenarios (Fig. 4D; supplementary Fig. S18). However, in scenarios where the recipient population size is reduced to 10% and 20% of the donor population size, the frequency of introgressed ancestries is negatively correlated with the level of recombination rates. These analyses suggest that the population size of the recipient population influences the correlation between the frequency of gene flow and the recombination rate. Thus, prolonged small population size of WL resulting from selective breeding may lead to the negative correlation between recombination rate and introgression frequency.

While adaptive introgression can occur, regions introgressed from divergent populations often introduce genetic incompatibilities that can reduce fitness, particularly in domestic species. Consistent with this, we observed a significant negative correlation between the frequency of deleterious mutations and the recombination rate (supplementary Figure S19). Our result reveals how deleterious mutations accumulate during the process of domestication and supports the “cost of domestication” hypothesis that domestication tends to induce the accumulation of deleterious in regions of low recombination (Lu et al. 2006).

## DISCUSSION

In this study, we employed population genomic approaches to construct high-resolution recombination maps of local and commercial chickens, as well as their wild relatives. Compared with earlier linkage-based maps (Groenen et al. 2009), our maps provide much finer resolution and enable more detailed comparative analyses. Despite the absence of the recombination-determining gene PRDM9, our results demonstrate that recombination landscapes in chickens are highly dynamic, with significant breed-specific variations. Notably, both commercial and local domestic chickens exhibit differentiation in effective recombination rates compared to their wild progenitor. This supports the idea that the rapid evolution of effective recombination is a common feature of domestication (Fuentes et al. 2022; Rees and Dale 1974; Lu et al. 2006).

Beyond the overall landscape, we identified significant variations in recombination hotspots across different populations, even those that diverged very recently. Most hotspots were population-specific, with little overlap with Red Junglefowl and among different populations. This rapid turnover is consistent with findings in other taxa, including sticklebacks, rice, chimpanzees, horses, mice and tomatoes. For example, a study of three-spined sticklebacks showed that the recombination landscapes of populations that diverged recently are highly divergent, with only around 15% of hotspots shared between different species. In rice, approximately 80% of hotspots were unique to each subspecies. These findings suggest that hotspots are largely short-lived and shaped by demographic history, genomic background and selection. However, it should be noted that the greater number of hotspots observed in domestic populations may partly reflect reduced recombination in surrounding genomic regions rather than entirely new hotspots emerging through de novo evolution.

Our analysis reveals a fundamental and robust inverse relationship between the location of selective sweeps and the local recombination rate, indicating that regions of low recombination are significantly enriched for the signatures of positive selection. We demonstrate that the strongest selective signals are confined to coldspots while noise is more randomly distributed, confirming that this pattern is not a methodological artifact but a genuine biological phenomenon. It is driven by the principles of Hill-Robertson interference, where low recombination facilitates the fixation of beneficial mutations and creates a detectable selective sweep signature. Similar findings have also been reported in horse breeds (Beeson, Mickelson, and McCue 2019), tomatoes (Fuentes et al. 2022) and apes (Li et al. 2019). For example, previous study in tomato demonstrated that the majority of selective sweeps were outside the recombination hotspot (Fuentes et al. 2022). In great apes, the majority of the recombination coldspots also largely overlapped with selective sweeps (Li et al. 2019). Therefore, domestication likely leads to the loss of the hotspot and/or the emergence of the coldspot, highlighting a specific way in which domestication may result in a decreased effective recombination rate (Fuentes et al. 2022). These findings suggest a close link between recombination remodeling and adaptation. The selective breeding of both domestic animals and crops typically aims to achieve phenotypic uniformity (Sae-Lim et al. 2017; Makumburage and Stapleton 2011). This practice involves rigorous selection to eliminate non-uniform traits and promote consistency, particularly in commercial breeds. For example, egg-laying poultry breeds such as WL have undergone intensive selection for traits such as exceptional egg production, efficient feed conversion, large egg size and early maturity over the last few decades. Although deleterious mutations tend to accumulate in regions with low recombination due to hitchhiking, mutations in a recombination coldspot benefits haplotype preservation, particularly when controlling for an ideal trait. However, the question of whether the emergence or disappearance of recombination hotspots and coldspots is associated with maintaining the uniformity and stability of phenotypic traits in domestic breeds during selective breeding remains unanswered.

Together, our results highlight profound dynamic recombination landscapes during the process of domestication and inbreeding. These findings contribute to a broader understanding of the genomic landscape of domestication, which is profoundly shaped by the interplay between recombination and selection. We acknowledge the potential limitations of estimating recombination from polymorphism data. Although we used medium-to high-coverage genomes to minimize error, complementary approaches, such as pedigree-based recombination mapping or sperm typing in crossbred populations, will be essential to validate and refine these patterns.

## MATERIALS AND METHODS

### Sampling and genome sequencing

Blood samples were obtained from 57 individuals, comprising 17 from the GGS, 20 from the WL, and 20 from the JNBR, and were stored in a freezer before DNA extraction. The GGS samples were collected earlier from Ruili in Yunnan province of China, while the WL samples were sourced from the farm of China Agricultural University. The JNBR samples were acquired from Shandong Hundred-Day Chicken Poultry Breeding Co., Ltd. DNA extraction was carried out using the phenol-chloroform extraction method. Paired-end DNA sequencing libraries were prepared, and sequencing was conducted on the Illumina NovaSeq 6000 platform, generating a total of 350.74 gigabytes of raw data. Detailed information regarding the sequencing data is available in supplementary Table S1. Additionally, genome re-sequencing data for 104 individuals, including 20 LX, 20 NX, 20 CH, 10 YOU, 10 SK, 20 GGM, 3 GGS along with a green junglefowl (*Gallus varius*; GV) genome used as an outgroup, were retrieved from the NCBI SRA database and ChickenSD (http://bigd.big.ac.cn/chickensd/; details could be found in supplementary Table S1).

### Sequencing alignment and variant calling

We removed the low-quality bases and reads using the trimmomatic v0.39 tool (Bolger, Lohse, and Usadel 2014) with parameters: “PE -threads 10 LEADING:3 TRAILING:3 SLIDINGWINDOW:4:15 MINLEN:36”. After qualification, all cleaned data were mapped to the chicken reference genome (GRCg7b) using default parameters except “-t 8 -M” option. We conducted processing on the BAM alignment, including coordinate sorting, marking duplicated reads, local realignment, and recalibrating base quality scores, utilizing the Picard (version 3.1.1; https://broadinstitute.github.io/picard/) and GATK (version 4.0; https://gatk.broadinstitute.org/hc/en-us) packages (McKenna et al. 2010). The average depth of coverage and the mapping ratio were provided in supplementary Table S1. We called SNPs following the GATK best practices pipeline (McKenna et al. 2010) using tools HaplotypeCaller and GenotypeGVCFs. We filtered out low-quality variants using the following criteria: “QD < 2.0 || FS > 60.0 || MQ < 40.0 || SOR > 3.0 || MQRankSum < −12.5 || ReadPosRankSum < −8.0”. As a result, 32,882,582 SNPs were obtained after quality filtration. To ensure the robustness of the dataset for each population, variants with minor allele frequencies below 0.05 and missing genotypes exceeding 20% in each population were meticulously excluded.

To further examine our result, we expanded our analysis to include whole-genome genotyping of ducks, pigs, goat, and sheep. Genotyping data for these species were sourced from previous studies (supplementary Table S3). Within each of these four species, there are four distinct breeds, each consisting of about 10 individuals.

### Population structure and phylogeny analysis

We meticulously examined the population structure across the entire genome for both GGS and domestic chicken breeds. We actively filtered SNPs based on linkage disequilibrium (LD) pruning using PLINK v1.9 (Purcell et al. 2007), employing the arguments: “-indep-pairwise 50 5 0.2”. Importantly, we deliberately excluded the sex chromosomes from our analysis. Leveraging this data, we constructed a maximum likelihood tree using the iqtree tool (Chernomor, von Haeseler, and Minh 2016), with parameters “-m TEST -st DNA -bb 1000 -nt AUTO”. Subsequently, we calculated the population LD decay using PopLDdecay (Zhang et al. 2019) and visually presented the LD decay profiles for all groups with default parameters.

The demographic history of each breed was inferred using SMC++ (Terhorst, Kamm, and Song 2017). We first generated genomic regions with uniquely mapped reads (“masked file”) using the tools in SNPable toolkit (http://lh3lh3.users.sourceforge.net/snpable.shtml). For each population, input files were prepared using the ‘vcf2smc’ tool (https://github.com/popgenmethods/smcpp) and the population size history was estimated using the ‘estimate’ function. For chicken, scaling was performed using a generation time of 1 year and a mutation rate of 1.91e-9 substitutions per site per year (Nam et al. 2010).

### Genomic summary statistics and selective sweep analysis

We performed a comprehensive analysis of genetic features within the genome of each breed and explored their correlation with recombination rate by calculating various summary statistics. For an in-depth examination of gene density and GC content across the chicken genome, we evaluated the proportion of gene regions in 2 Mb non-overlapping sliding windows based on the gene model annotation file (gff format). To determine the genetic diversity within the six chicken breeds, we calculated the nucleotide diversity (pi) for each breed using vcftools with the parameters “--window-pi 50000” (Danecek et al. 2011).

We used selscan version 1.3.0 (Szpiech and Hernandez 2014) to identify genomic regions with signals of selection in each breed. Genome-wide nSL scores were normalized with selscan’s companion program norm from selscan using default parameters. We partitioned the genome into non-overlapping 50 kb regions and calculated the fraction of sites with |nSL| > 2 in each window. We then selected the windows with the top 1%, 5%, and 10% of the fraction of sites with high scores as potential sweeps, which were used for further detailed analysis.

We also used, as a complementary method, the locus-specific branch length (LSBL) statistic (Shriver et al. 2004) to scan the signal of selection, which was calculated using a sliding window approach with a window of 50kb and a step of 25kb.

### Inference of locally introgressed ancestries

Using high-quality sequencing datasets comprising 20 samples from WL, 20 from CH, and 20 from GGM (supplementary Table S1), we identified segments in the WL population that were introgressed by GGM. First, we calculated “fd” (Martin, Davey, and Jiggins 2015), a metric based on the calculation of ABBA-BABA sites in the four-taxon topology (((P1, P2), P3), O), to detect and quantify bidirectional introgression at specific loci. In our calculation, we designated the CH, WL, and GGM populations as P1, P2, and P3, respectively, with a green junglefowl serving as the outgroup. We calculated “fd” values using a 100 kb sliding window with 20 kb increments, utilizing the command: “ABBABABAwindows.py -w 100000 -m 10 -s 20000 ChrN.vcf.geno.gz -o fd.csv.gz -f phased --minData 0.5 --writeFailedWindow - P1 CH -P2 WL -P3 GGM -O GV --popsFile pop.info”. Windows with fdM>0 and the number of informative sites ≥ 10 were retained as outlier loci.

Finally, we utilized Loter to infer local ancestry, a method demonstrated to be robust in detecting more ancient admixtures, to obtain admixed ancestry in WL from GGM (Dias-Alves, Mairal, and Blum 2018). We specified the GGM and CH as reference panels, with WL being the admixed population with the command “loter_cli -a WL.ChrN.vcf -r ChaHua.ChrN.vc f GGM.ChrN.vcf -f vcf -n 6 -v -pc -o WL.ChrN.CH-GGM.Loter.txt”. Loter determines the most likely ancestral origin for each fragment using allele frequencies from the reference and selected populations. Introgression percentages from GGM into haploid genomes were calculated, and overlapping introgressed regions from the same source were merged. Frequencies were then calculated for each selected fragment across the WL.

### Recombination rate estimation

We used RelERNN (v1.0.0) to estimate population-scale recombination rates for each chromosome in six chicken populations (Adrion, Galloway, and Kern 2020). RelERNN uses recurrent neural networks, and takes columns from a genotype alignment, which are then modelled as a sequence across the genome using a recurrent neural network. We trained the datasets with parameters based on the developers’ recommendations under both the equilibrium demographic scenario and the SMC++ inferred demographic scenario (“--demographicHistory” option).

To evaluate the robustness of our results across different parameters, we examined the impact of different gradient parameters and identified the “--nEpochs” parameter as the most influential. Consequently, we performed several simulated training runs under the equilibrium demographic scenario, varying the “--nEpochs” parameter from 100 to 2000 iterations (100, 200, 500, 1000, and 2000). Remarkably, using more than 500 training runs (--nEpochs 500) resulted in a correlation coefficient greater than 0.9. Despite additional training efforts (e.g., 1000 and 2000 runs), the correlation showed only marginal improvement, suggesting near saturation. Considering the computational resources used, we chose 1000 training runs for the subsequent analyses.

To explore the effect of sample size on the estimation accuracy, we used RelERNN to estimate the recombination rate under both the equilibrium demographic scenario using 5, 10, and 20 samples with the settings “--nEpochs 1000, MU=1.91e-9, --nTrain 100000, --nVali 1000, --nTest 1000, and --nValSteps 20”. We measured the correlation of each estimate using Spearman’s correlation. Notably, our results showed a high degree of consistency in the estimates when the sample size was 10 or greater.

Therefore, based on the results of the above tests, we estimated recombination rates for GGS and each chicken breed, as well as combining all samples (including both GGS and domestic chickens) using parameters “--nEpochs 1000, MU=1.91e-9, -- nTrain 100000, --nVali 1000, --nTest 1000, and --nValSteps 20” with 20 samples per chicken breed. We then standardized them into non-overlapping 50-kb windows. Sex chromosomes and dot chromosomes (Chr16, 29-32,34, and 39) were excluded from the analyses because of incomplete assembly and the large proportion of missing rate.

The similarity of recombination distribution among chicken breeds was assessed using Pearson’s correlation, both for whole genome comparisons (1 Mb intervals) and individual chromosomes.

### Identify recombination hotspots and coldspots

Recombination hotspots and coldspots were identified according to methods outlined in a prior study (Palahi et al. 2023). First, we divided the recombination map into non-overlapping 1 kb windows. Hotspots were characterized as regions exhibiting a recombination rate at least 2 times higher than the regional “background” rate, determined as the average rate for the 50 kb focal window and its two flanking windows, in total 150 kb. Conversely, coldspots were regions with a recombination rate at least 10 times lower than the regional background rate.

To evaluate overlap with selective sweeps, gene flow, etc., and assess chance outcomes, we generated 100/10,000 sets of random, size-matched genomic segments to mimic detected hotspots/coldspots using “shuf -n”. These simulated regions were termed “randomspots,” and overlap was assessed using bedtools intersect. Fisher’s Exact tests identified genomic features with differential enrichment between hotspots/coldspots.

Hotspots were further analyzed for proximity to promoters, coding sequences (CDS), and intergenic regions. Overlap with previously published double-strand break (DSB) hotspots identified by ChIP-seq was also examined using the intersect function in the bedtools software. Additionally, we analyzed hotspot/coldspot overlap within and between subspecies by examining population-specific sets using bedtools intersect.

### Identify recombination hotspot determining motif

To determine whether there are any critical motifs involved in chicken recombination hotspots, we employed RHSNet, a cutting-edge tool rooted in deep learning and signal processing (Li, Chen, et al. 2022). RHSNet is tailored to predict significant motifs contributing to recombination hotspots specifically in chickens. Our methodology commenced with constructing a comprehensive dataset for 5-fold cross-validation and motif extraction, incorporating datasets such as the Icelandic Human dataset (Halldorsson et al. 2019), the HapMap II dataset (Frazer et al. 2007), the Sperm dataset (Bell et al. 2020), hotspots and coldspots derived from paternal and maternal maps of the Icelandic Human dataset (Halldorsson et al. 2019), along with the mouse dataset (Lange et al. 2016). Then, we conducted training sessions, including Baseline CNN training, Equivalent CNN training, RHSNet training, RHSNet-Chip training with consideration for sexual differences, and Baseline CNN training on the 1000 Genome Dataset. For motif extraction, we employed the Python script “motif_extractor.py” from RHSNet, facilitating the identification and analysis of key motifs within the chicken recombination hotspots.

### Identification of deleterious mutations

To predict the functional impact of missense mutations, we applied PROVEAN (Choi and Chan 2015) to compare query sequences against the NCBI non-redundant protein database. We identified mutations with PROVEAN scores of ≤-2.5, which indicate deleterious effects.

### Simulation analysis

To explore the impact of effective population size on the correlation between recombination and gene flow, we conducted a simulation analysis. Our model is grounded in a four-population topology (((P1, P2), P3), O) reflecting the phylogeny and splitting of GV, GGM, GGS, and WL. In the model, P4 diverged from P1-P2-P3 around 4 million years ago, P3 diverged from P1-P2 approximately 80,000 years ago (Wang et al. 2020; Wang et al. 2021), and P1 diverged from P2 around 10,000 years ago, and divergent P3 population (donor) contributed 10% of the gene flow to the recipient P1 population. P3 and P2 maintain a consistent population size of N0, while P1 exhibits different population sizes in various models. We tested five models in which P1 has population sizes of 10*N0, 2*N0, N0, 0.2*N0, and 0.1*N0, respectively, in 200 years (corresponding to the breeding history of WL). In each model, we used msprime (Baumdicker et al. 2022) to simulate 2, 40, 40, and 40 haplotypes with a length of 2Mb for O, P3, P2, and P1, respectively. Thirty replicates were performed for each simulation. Utilizing the simulated data, we used Loter to identify P3 introgressed segments in P1, and estimated the recombination rates using RelERNN. Subsequently, we measured the Pearson’s correlation between the frequency of introgressed ancestry and recombination rate. In our simulations, N0 was set to 10000, and the mutation rate (per site per year) was set to 1.91e-9.

## Supporting information

supplemental files will be used for the link to the file on the preprint site

## ACKNOWLEDGMENTS

This work was supported by the National Key Research and Development Program of China (2023YFF1001000) and the Yunnan Fundamental Research Projects (202301AW070012, 202401AV070007), Yunnan Province (202305AH340006) and the “Yunnan Revitalization Talent Support Program: High-end Foreign Expert Project and Young Talent Project (XDYC-QNRC-2022-0770)”. The National Natural Science Foundation of China, Talent Program of Chinese Academy of Sciences (CAS) and Animal Branch of the Germplasm Bank of Wild Species of CAS (the Large Research Infrastructure Funding) also supported this project. A.E. was supported by the CAS President’s International Fellowship Initiative (No. 2024VBA0001). We would like to thank Hong-Man Chen for his assistance in preparing the figures and Jeffrey R. Adrion for his support in running RelERNN.

## AUTHOR CONTRIBUTIONS

Ming-Shan Wang, Ya-Ping Zhang, and Jian-Lin Han conceived the project and designed the research. Zheng-Xi Liu and Ming-Shan Wang performed the analysis. Jian-Lin Han and Lu-Jiang Qu collected samples and prepared the DNA samples and contributed genome sequencing data. Ming-Shan Wang and Zheng-Xi Liu drafted the manuscript with input from all authors. Xue-Hai Ge, Kun Wang, and Chang-Rong Ge provided the pedigree data. Ming-Shan Wang, Ming Li, Ya-Ping Zhang, Si Si, Jian-Lin Han, Jian-Hai Chen, Chang Zhang, Li-Rong Hu, Min-Sheng Peng, Ting-Ting Yin, Ali Esmailizadeh, and Xue-Mei Lu revised the manuscript. All authors read and improved the final manuscript.

## DATA AVAILABILITY

The 57 whole genomes we generated were submitted to GSA (https://ngdc.cncb.ac.cn/gsa/) under the accession number PRJCA025697. Scripts and recombination estimations are available at https://github.com/Joyce121. compliance and ethics

## SUPPLEMENTARY DATA

Supplementary tables and figures are available at Supplementary information.

