## supplemental files will be used for the link to the file on the preprint site for "Rapid evolution of fine-scale recombination during domestication: a perspective from population genomics"

Supplementary Materials include Supplementary Figures S1-18 and Supplementary Tables 1-3.

**Supplementary Figures**

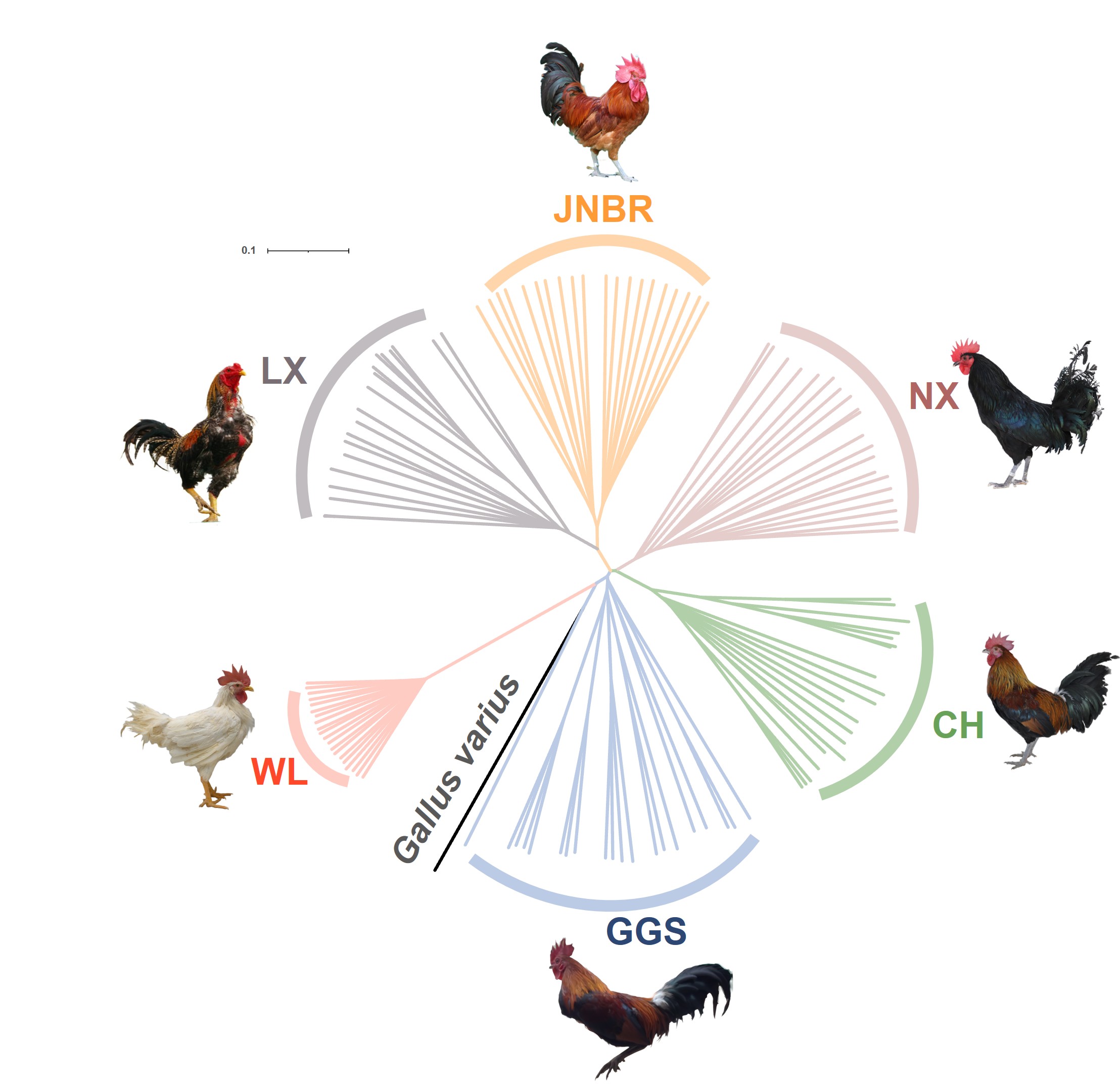

**Fig. S1.** Phylogenetic tree showing the relationship between domesticated chickens and GGS. The tree was constructed using IQ-TREE software based on the maximum likelihood method. The photos of WL, GGS, NX, CH, LX and JNBR were taken by the following people and we have their permission to use them. WL by Wendi Ma (University of Chinese Academy of Sciences), NX and CH by Tengfei Dou (Yunnan agricultural university), LX by Xiang Gao (Breeding Farm of Cockfighting Roosters, Shandong, China) and Xuxian Qu (Provincial Station of Animal Husbandry, Shandong, China) and JNBR by Xuxian Qu and Ke Qu (Adminstration of Animal Husbandry and Veterinary Service, Jining, Shandong, China), GGS by Chatmongkon Suwannapoom (School of Agriculture and Natural Resources, University of Phayao, Phayao, Thailand).

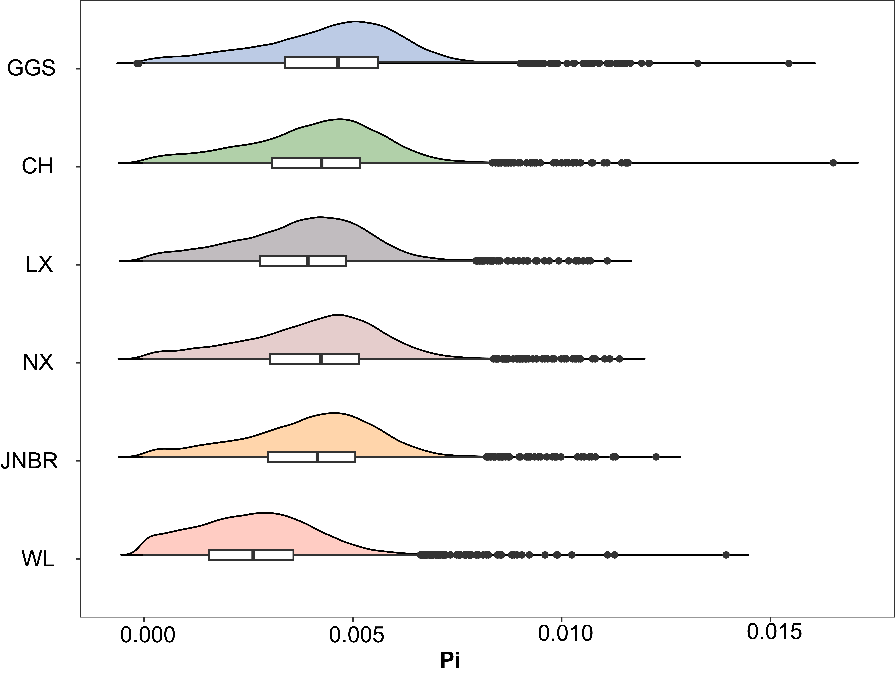

**Fig. S2.** Genetic diversity(π) for GGS and domestic chickens.

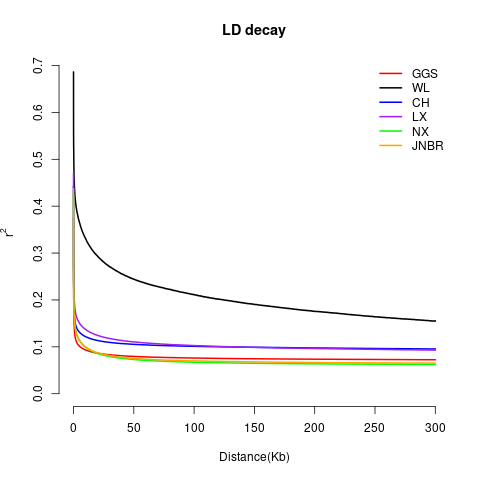

**Fig. S3.** The decay of linkage disequilibrium (LD) over genetic distance for GGS and domestic chickens.

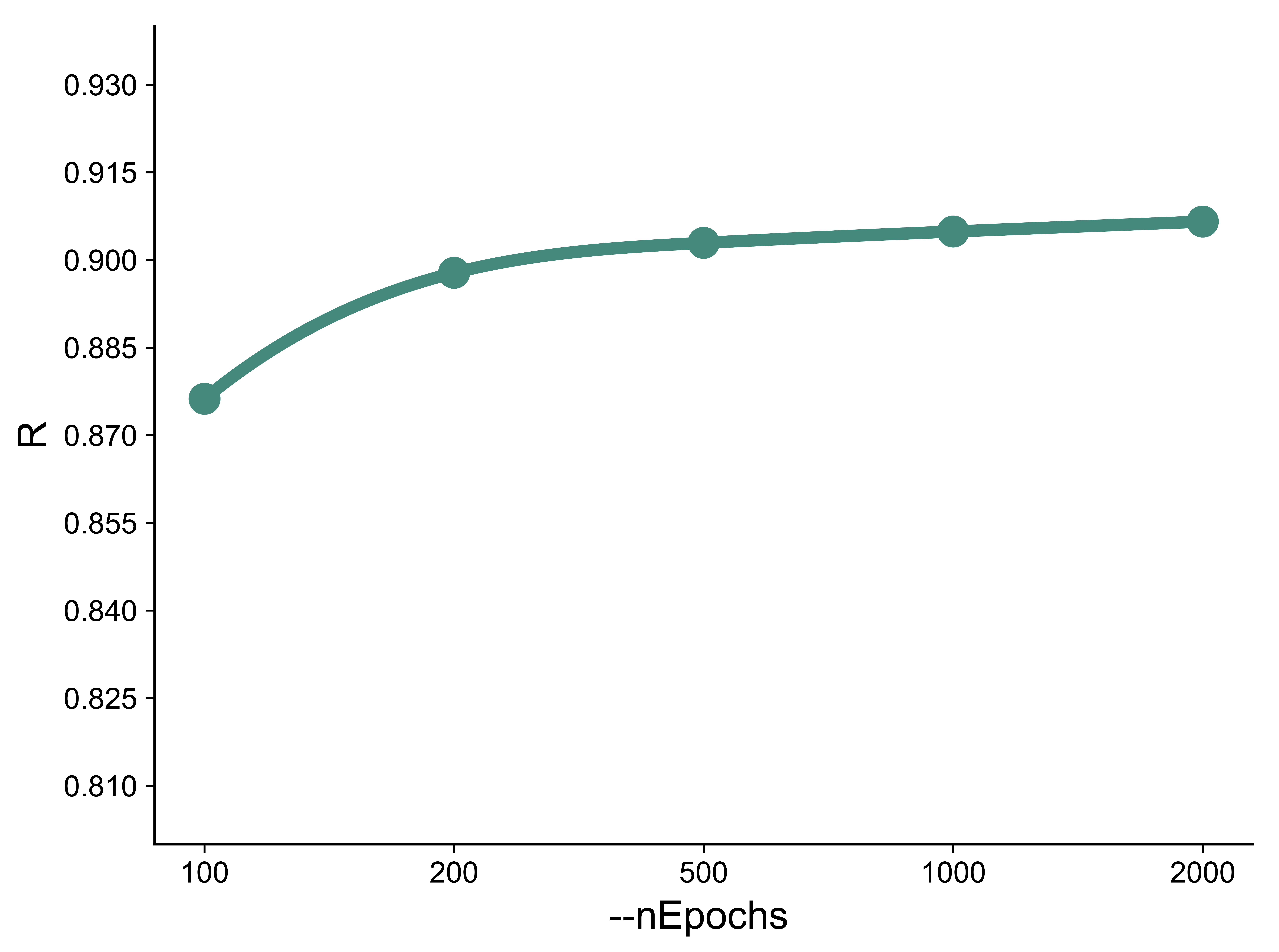

**Fig. S4.** Effect of training time on prediction correlation coefficients. An overall trend of the correlation coefficient from 500 to 2000 epochs is shown for comparison.

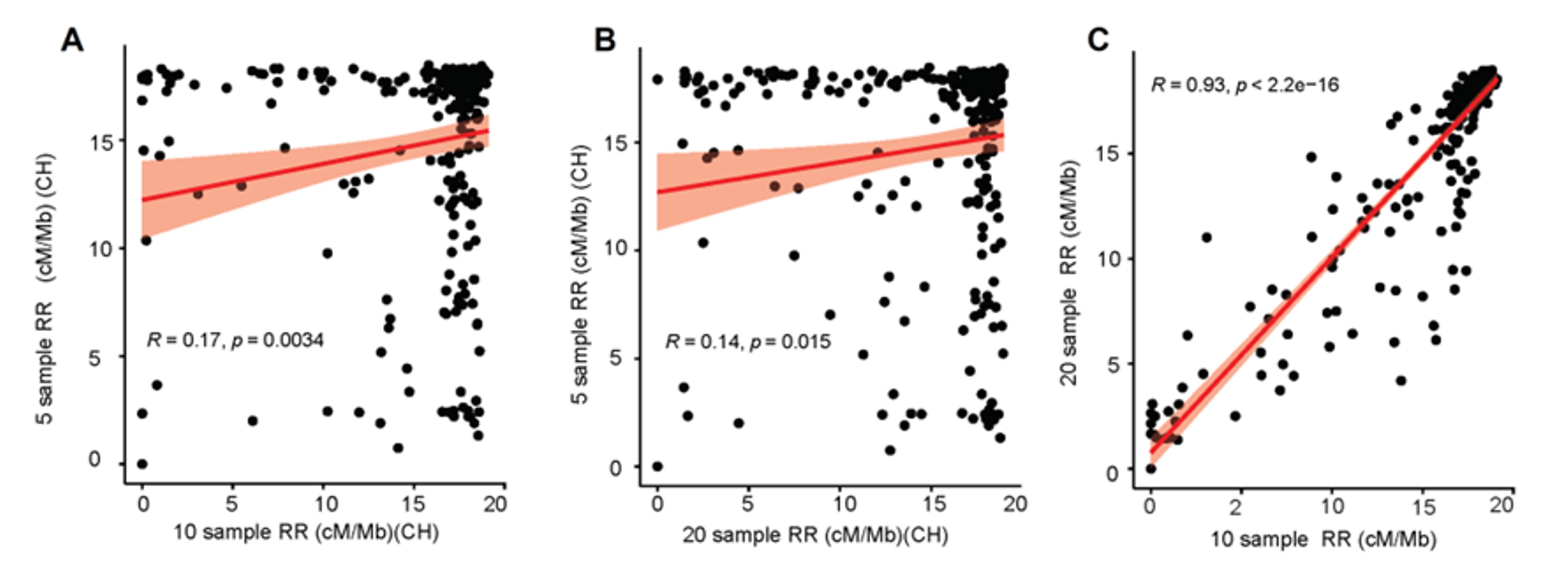

**Fig. S5.** Evaluation of correlation coefficients for recombination rates over different sample sizes. The correlation coefficients derived from the analysis of recombination rates corresponding to different sample sizes: subgroup (A) consisting of 5 individuals, subgroup (B) consisting of 10 individuals, and subgroup (C) consisting of 20 individuals.

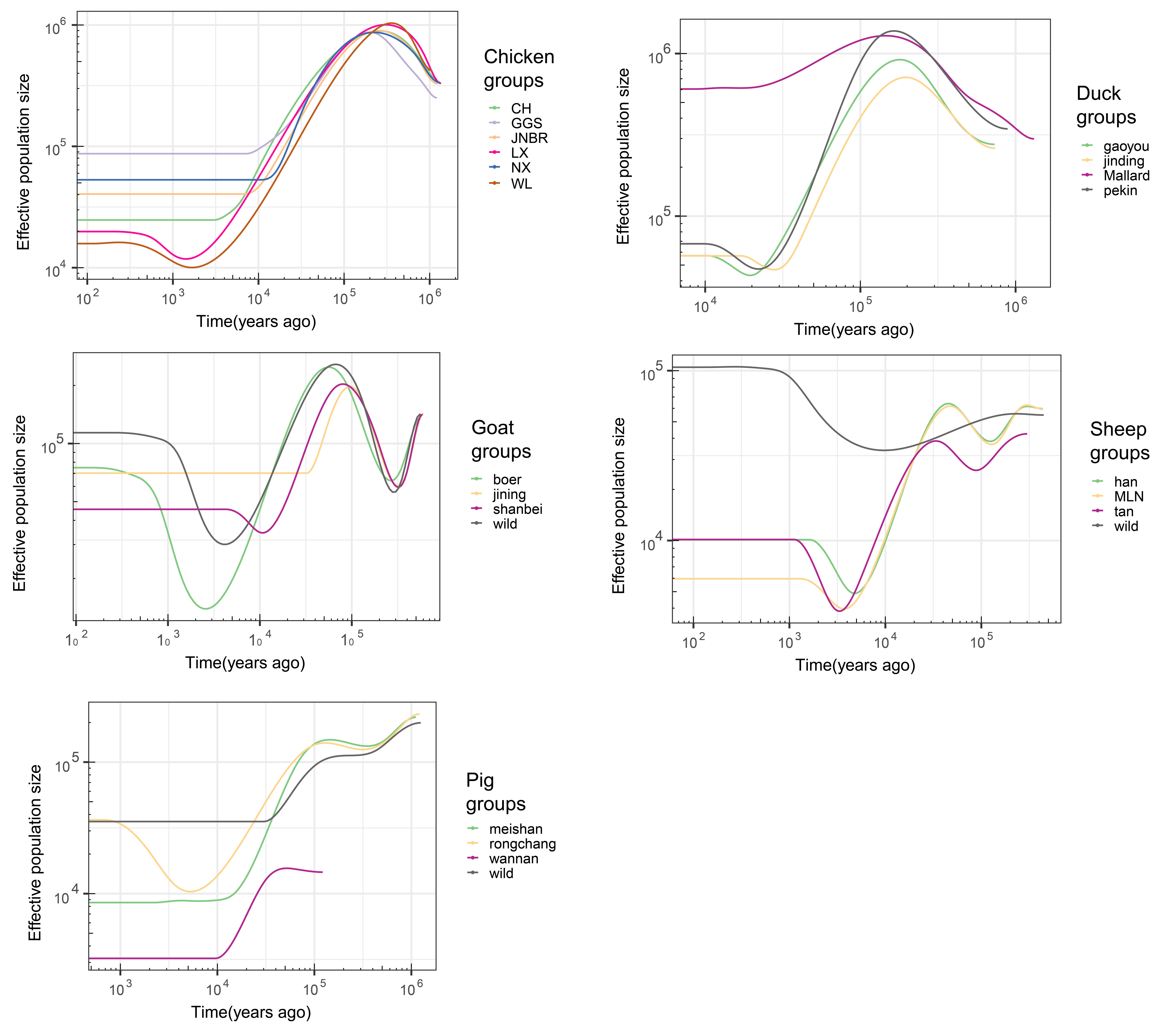

**Fig. S6.** The demographic history for each species (chicken, duck, goat, sheep and pig) is shown below. For chicken, the mutation rate was 1.91e-9 substitutions per site per year and the generation time was 1 year [1]. For duck, the mutation rate was 1.61e-9 substitutions per site per year and the generation time was 1 year [2]. For goat, it was 2 years and 4.32e-9 substitutions per site per year. For sheep, it was 3 years and 1.51e-8 substitutions per site per year [3]. The same was done for pigs, using a generation time of 3 years and a mutation rate of 3.6e-9 substitutions per site per year [4].

1. Nam K, Mugal C, Nabholz B, Schielzeth H, Wolf JB, Backström N, et al. Molecular evolution of genes in avian genomes. Genome Biol. 2010;11(6):R68.
2. Zhang Y, Chen Y, Zhen T, Huang Z, Chen C, Li X, et al. Analysis of the genetic diversity and origin of some Chinese domestic duck breeds. J Integr Agric. 2014;13(4):849–57
3. Zhuqing Zheng et al., The origin of domestication genes in goats.Sci. Adv.6,eaaz5216(2020).
4. Zhang M, Yang Q, Ai H, Huang L. Revisiting the Evolutionary History of Pigs via De Novo Mutation Rate Estimation in A Three-generation Pedigree. Genomics Proteomics Bioinformatics. 2022 Dec;20(6):1040-1052.

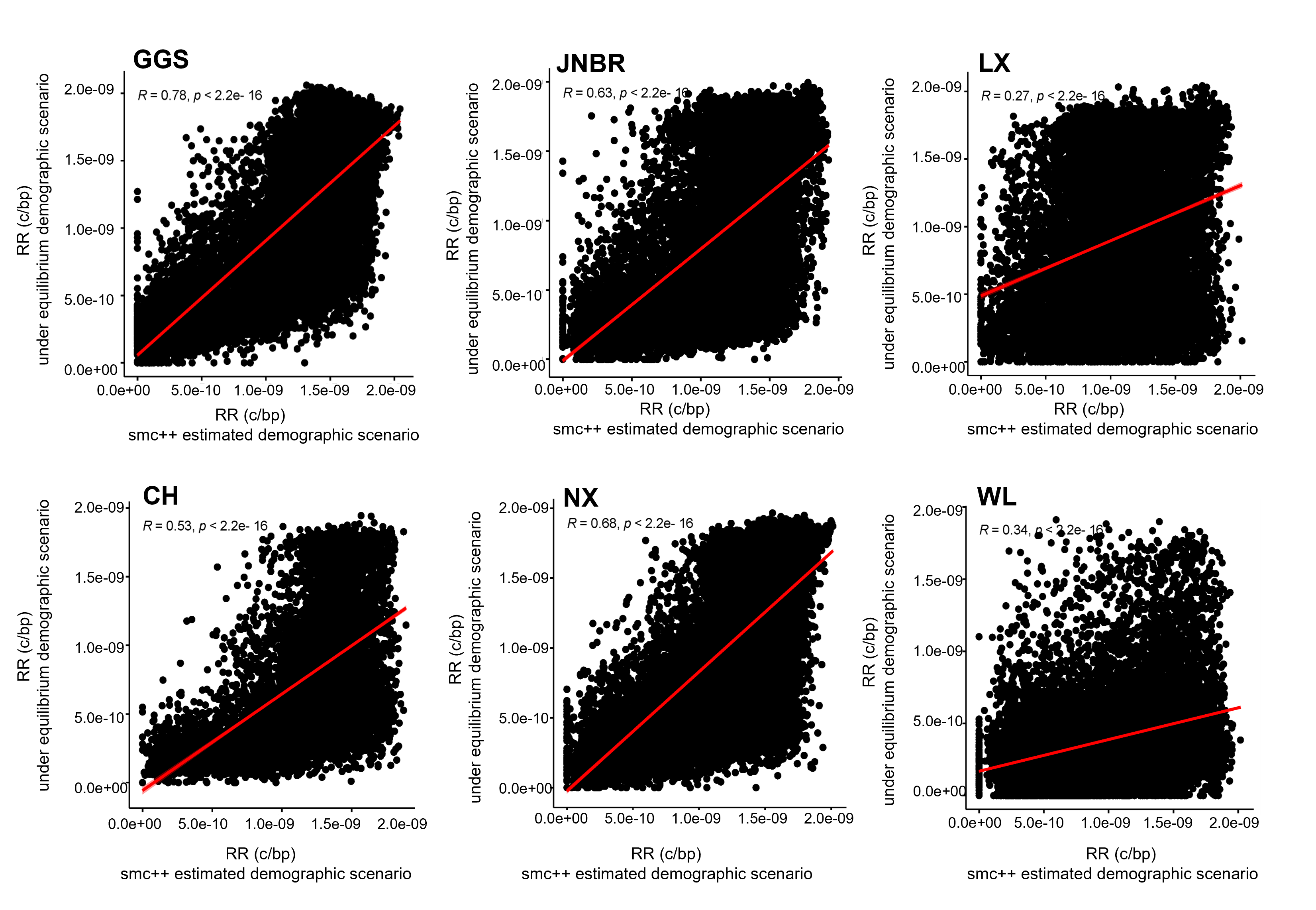

**Fig. S7**. Pearson's correlation of the recombination rate between using an equilibrium model and a demographic history model inferred by SMC++ of each population.

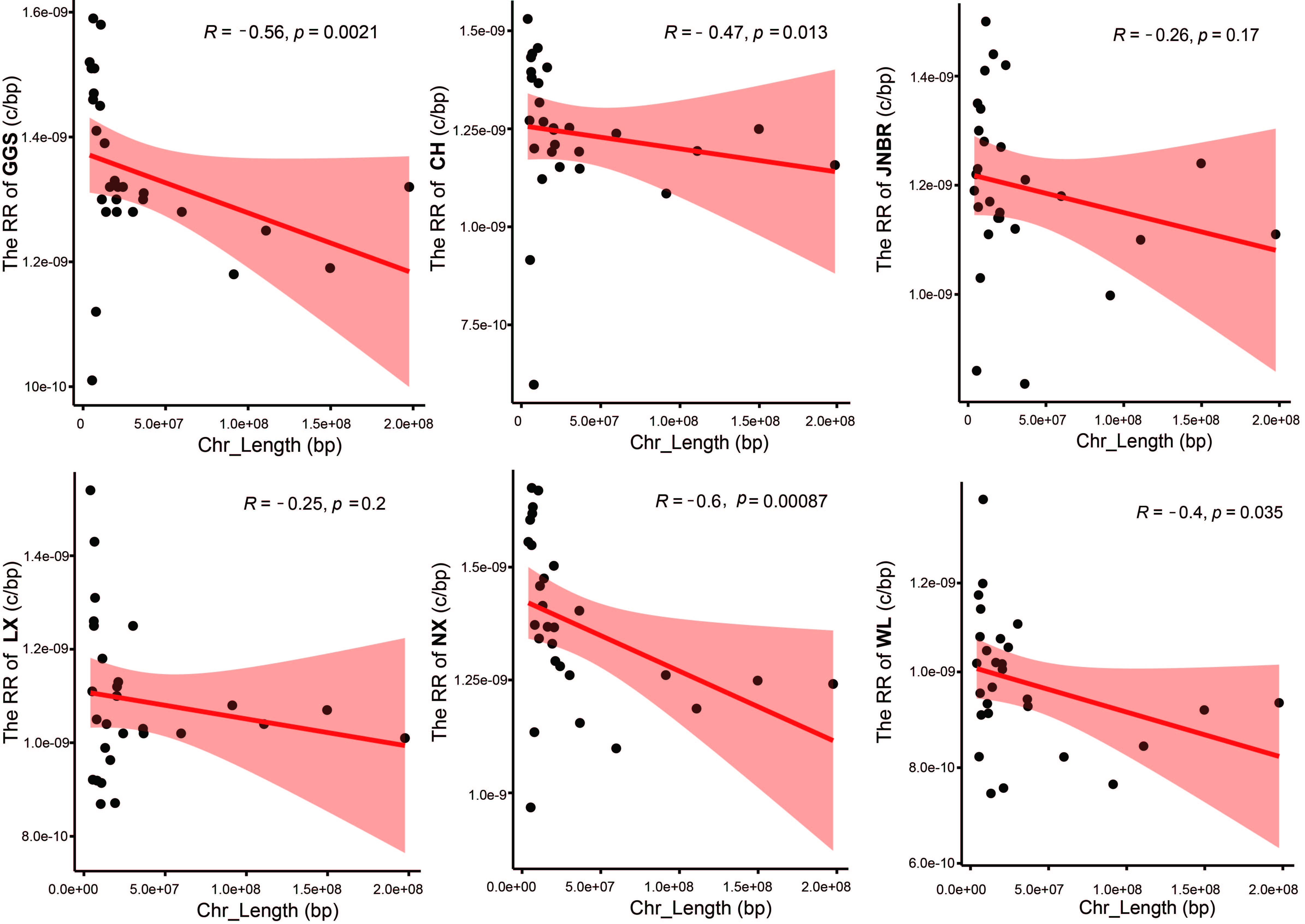

**Fig. S8.** Pearson's correlation between recombination rate and chromosome size in each chicken breed.

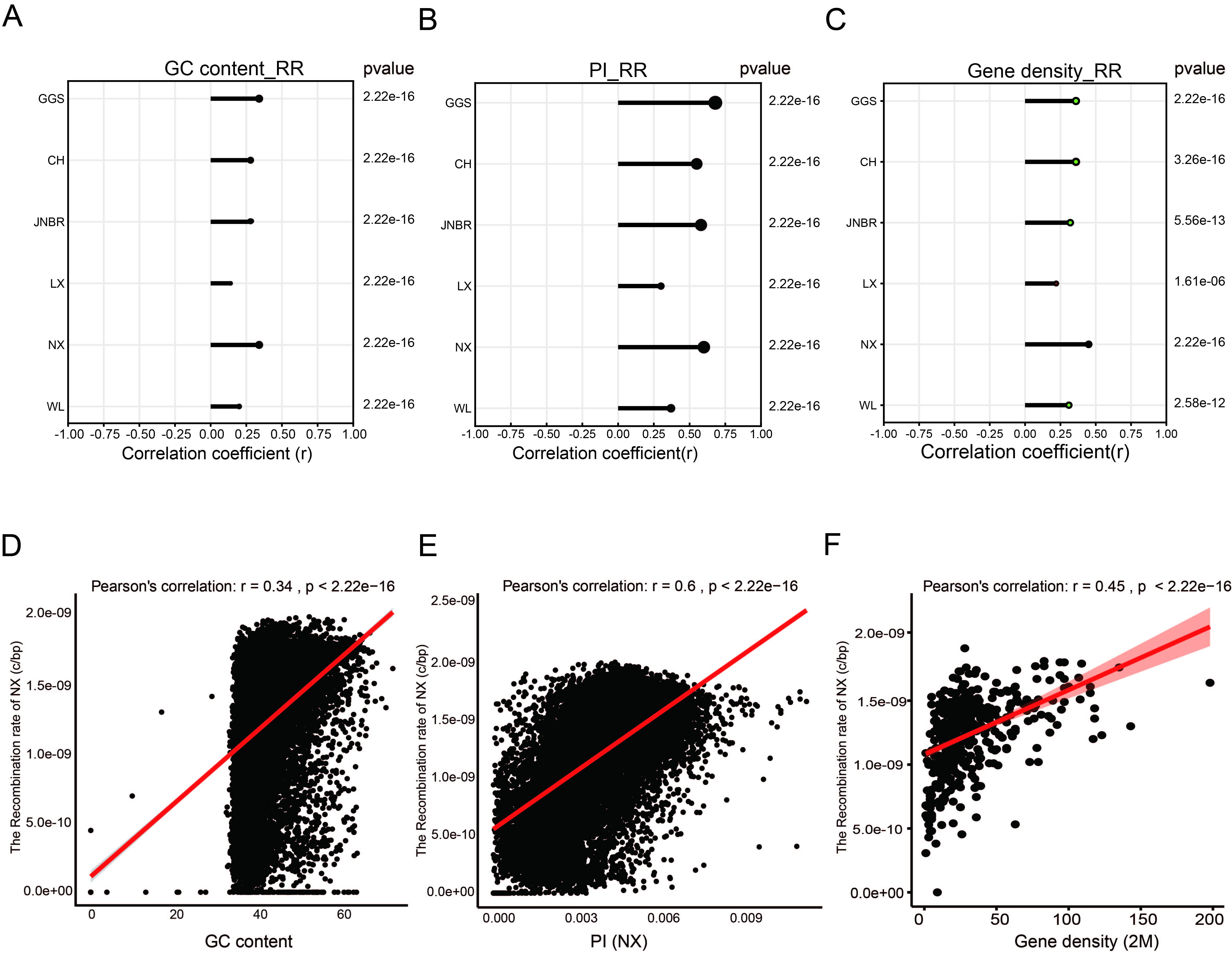

**Fig. S9.** Correlation coefficients for recombination rate and genomic characteristics in six breeds. (A) Pearson's correlation between recombination rate and GC content in each chicken breeds. (B) Pearson's correlation coefficient between recombination rate and genetic diversity across the breeds. (C) Pearson's correlation between recombination rate and gene density in each chicken breed. (D-F) Correlation of recombination rate with GC content, genetic diversity (Pi), and gene density respectively, illustrated with a single breed as an example.

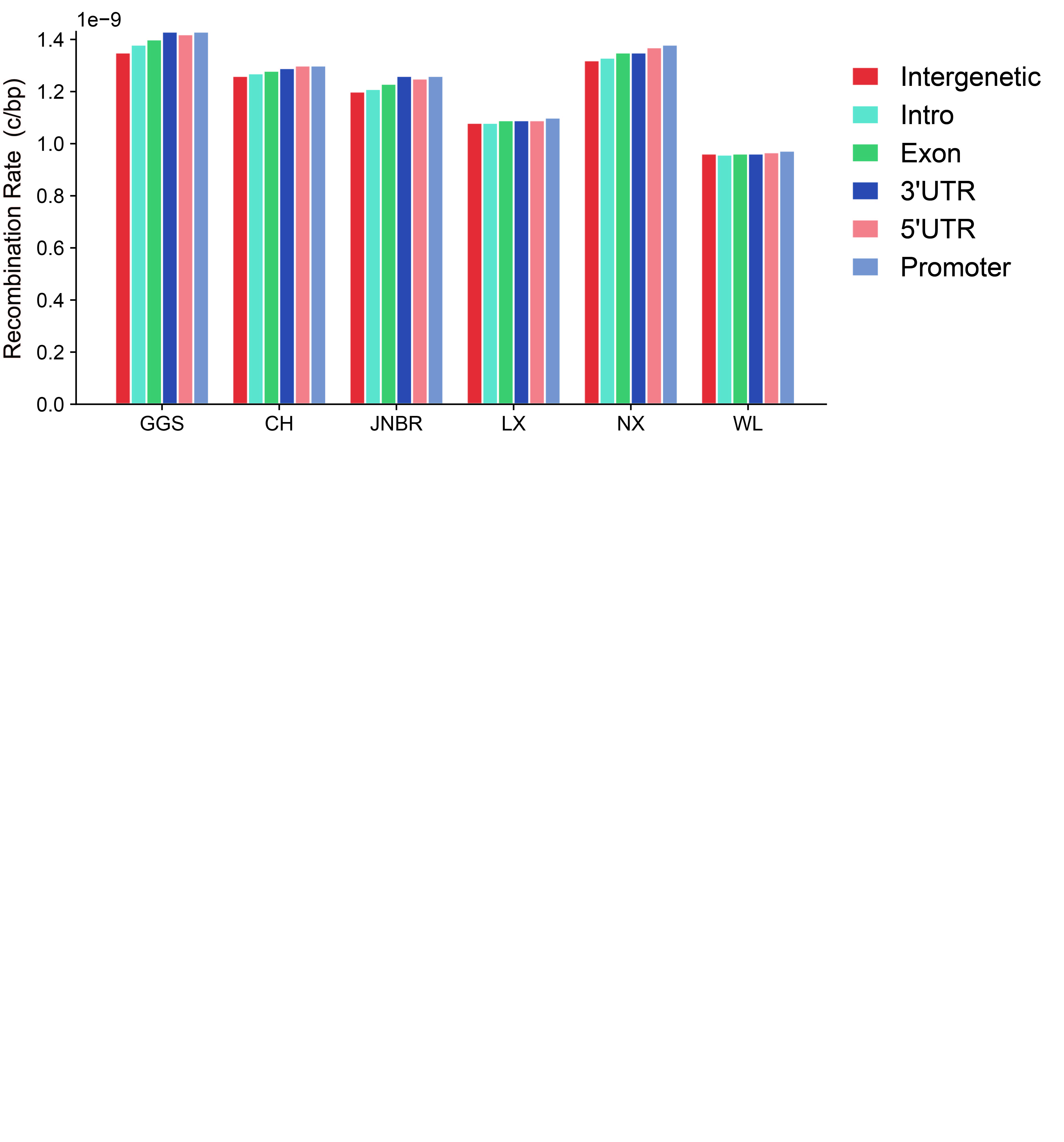

**Fig. S10.** Distribution of recombination rate in respective to intergenic, intron, exon, 3' untranslated region (3' UTR), 5' untranslated region (5' UTR), and promoter regions.

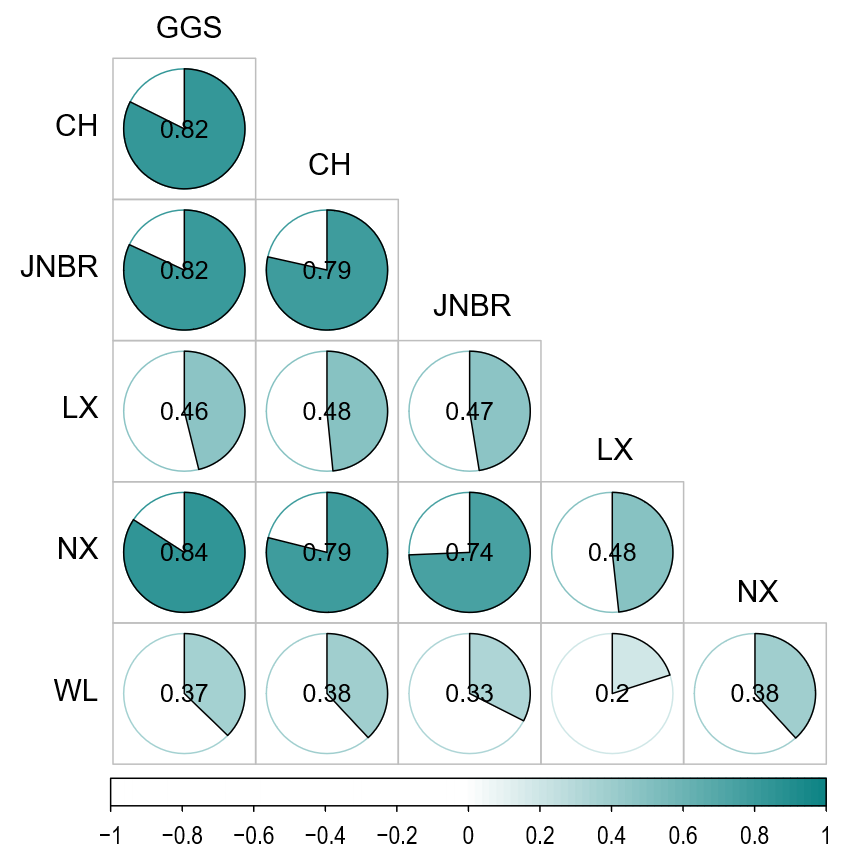

**Fig. S11.** The Pearson's rank correlation coefficients of recombination rates among six breeds.

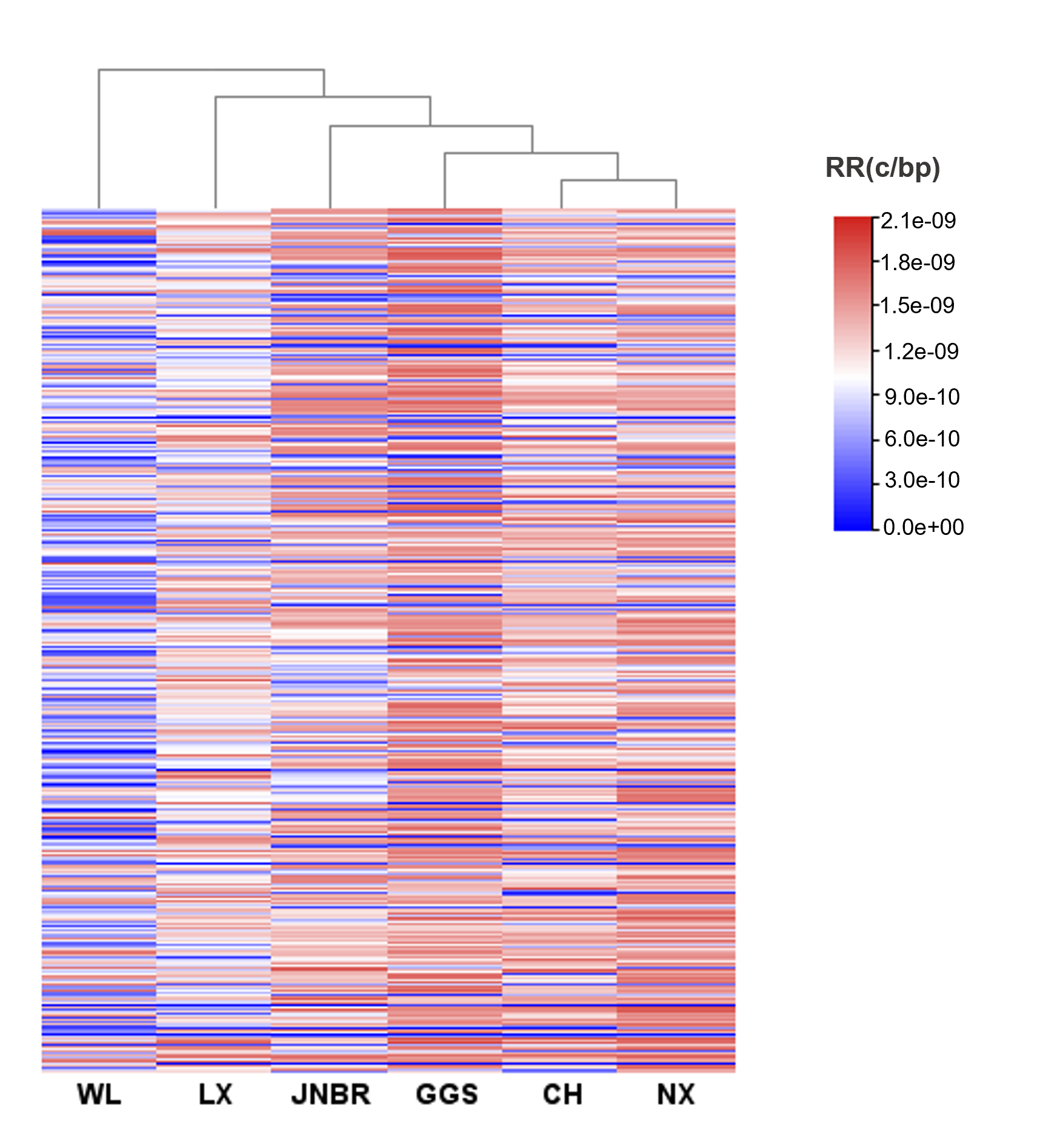

**Fig. S12.** Phylogenetic reconstruction of the studied populations based on recombination rate

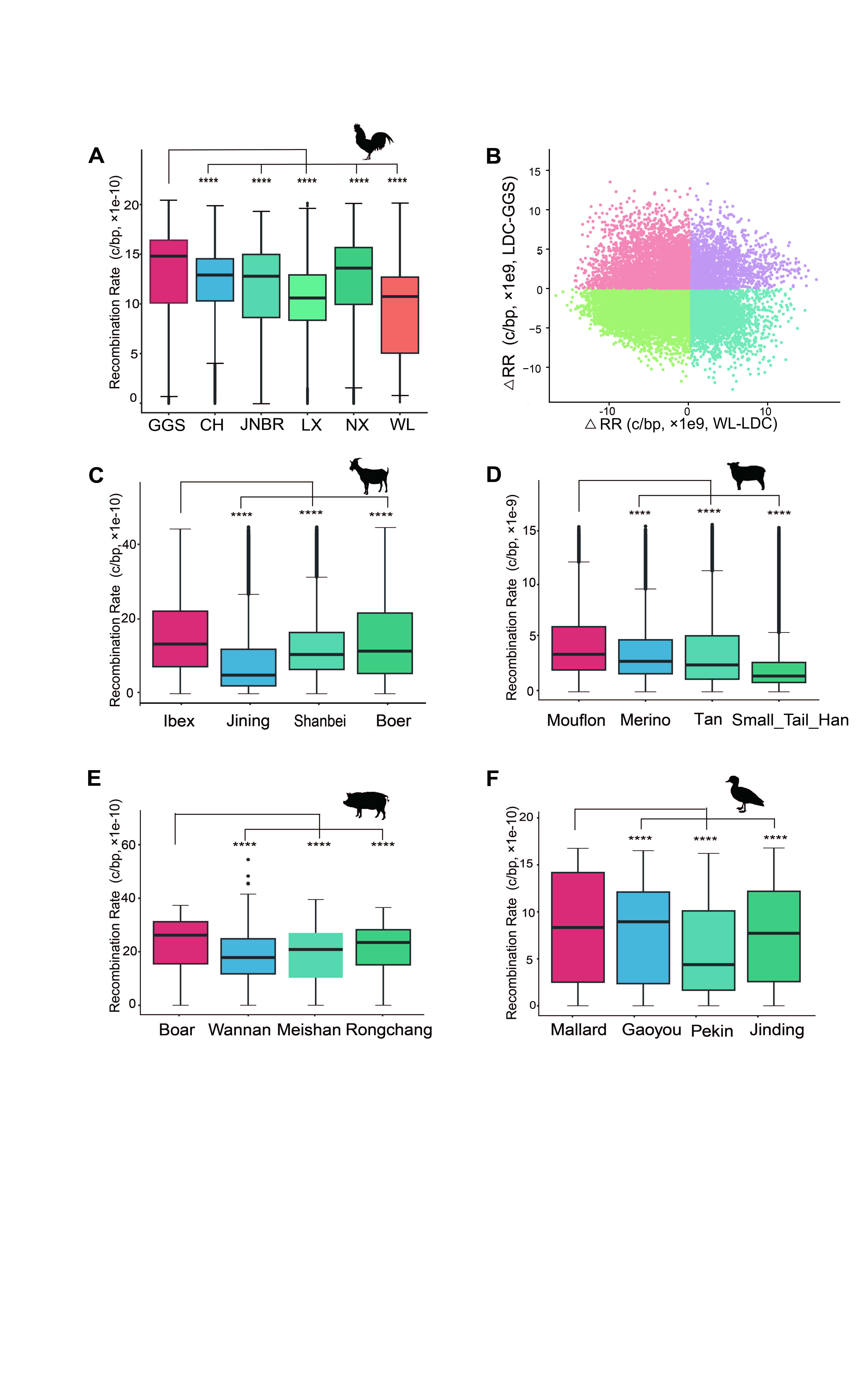

**Fig. S13.** Recombination landscape between domesticated species and their wild relatives. (A) Recombination rates of Red Jungle Fowl (*Gallus gallus spadiceus*, GGS) and domesticated chickens. (B) The level of recombination rate (RR) changes in LDC and WL compared to GGS. (C) Recombination rates of Bezoar ibex (*Capra aegagrus*) and domesticated goats. (D) Recombination rates of mouflon (*Ovis gmelini*) and domestic sheep. (E) Recombination rates of Asian wild boar (*Sus scrofa*) and domestic pigs. (F) Recombination rates of mallard duck (*Anas platyrhynchos*) and domestic ducks. In this figure, **** indicate *P* < 0.0001 by the Wilcoxon test.

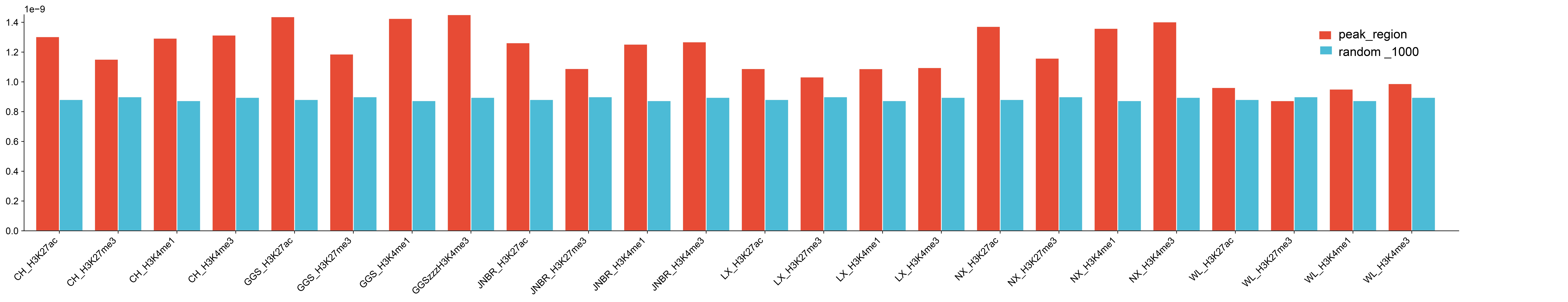

**Fig. S14.** Comparison of recombination rates between regions associated with four histone modifications (H3K27ac, H3K27me3, H3K4me1, and H3K4me3) and randomly selected regions (random_1000, n=1000) of equal length from the rest of the genome.

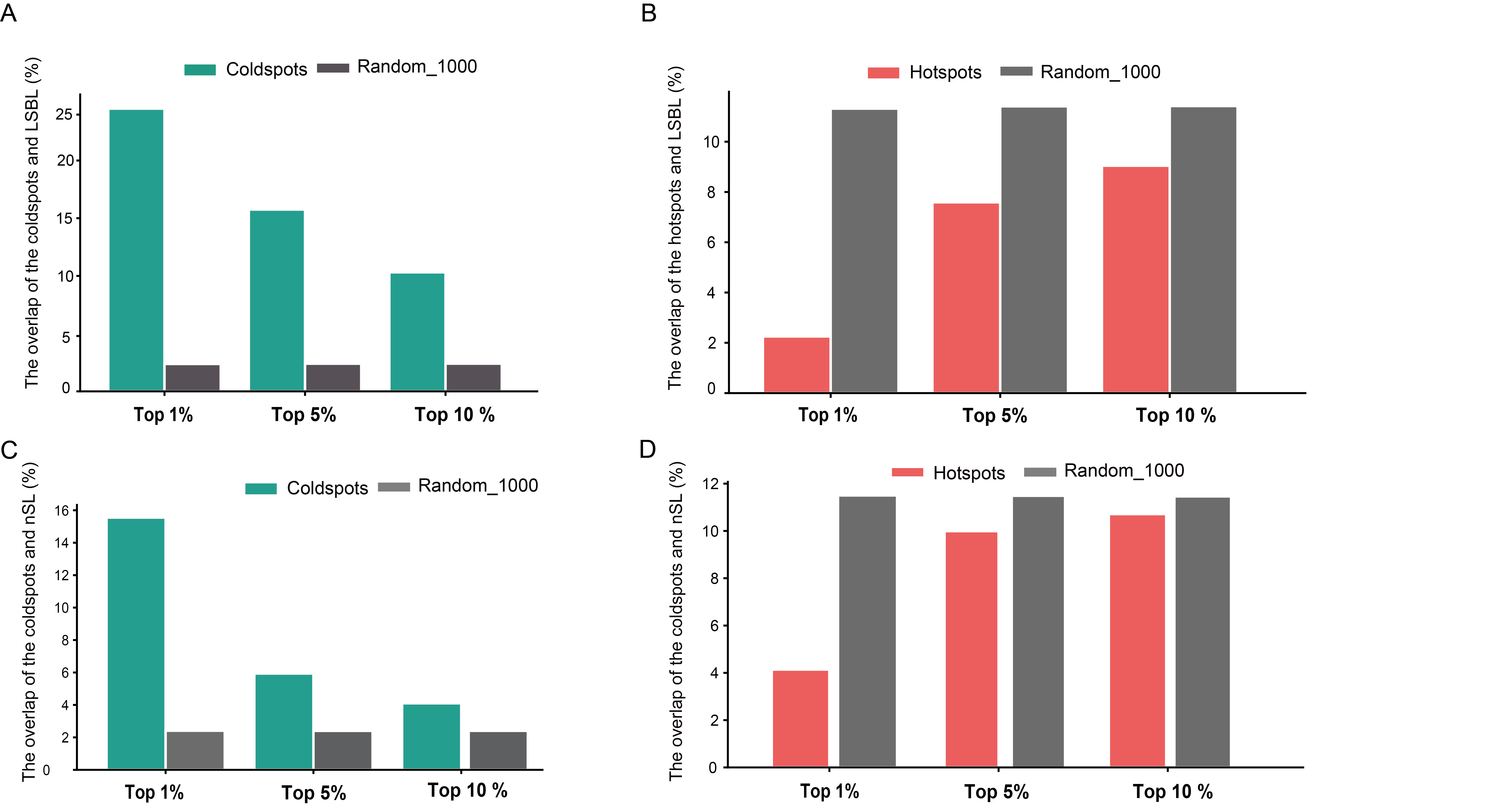

**Fig. S15.** The overlap between hotspots/coldspots and putative selective sweeps (identified by LSBL and nSL top 1%, top 5% and top 10%) of WL. (A) The overlap rate between recombination coldspots and putative selective sweeps (identified by LSBL) is higher than that of randomly selected regions(n=1000). (B) The overlap rate between recombination hotspots and putative selective sweeps (identified by LSBL) is lower than that of randomly selected regions(n=1000). (C) The overlap rate between recombination coldspots and putative selective sweeps (identified by nSL) is higher than that of randomly selected regions(n=1000). (D) The overlap rate between recombination hotspots and putative selective sweeps (identified by nSL) is lower than that of randomly selected regions(n=1000).

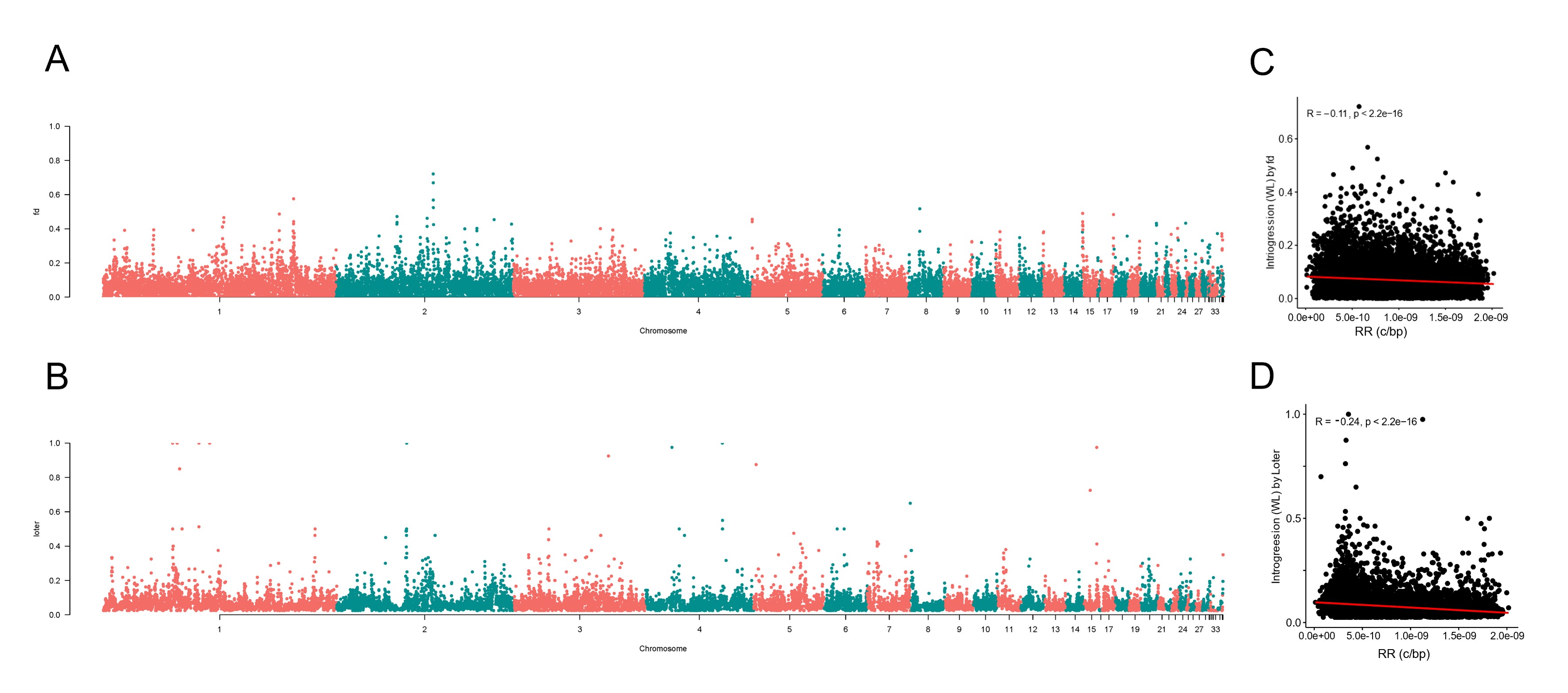

**Fig. S16.** Relationship between recombination rate and frequency of introgressed ancestry calculated by Fd (A, C), Loter (B, D). (A) Manhattan plots of introgressed ancestry identified by Fd. (B) Manhattan plots of introgressed ancestry identified by Loter. (C) Correlation coefficients between introgressed ancestry by Fd and recombination rate. (D) Correlation coefficients between introgressed ancestry by Loter and recombination rate.

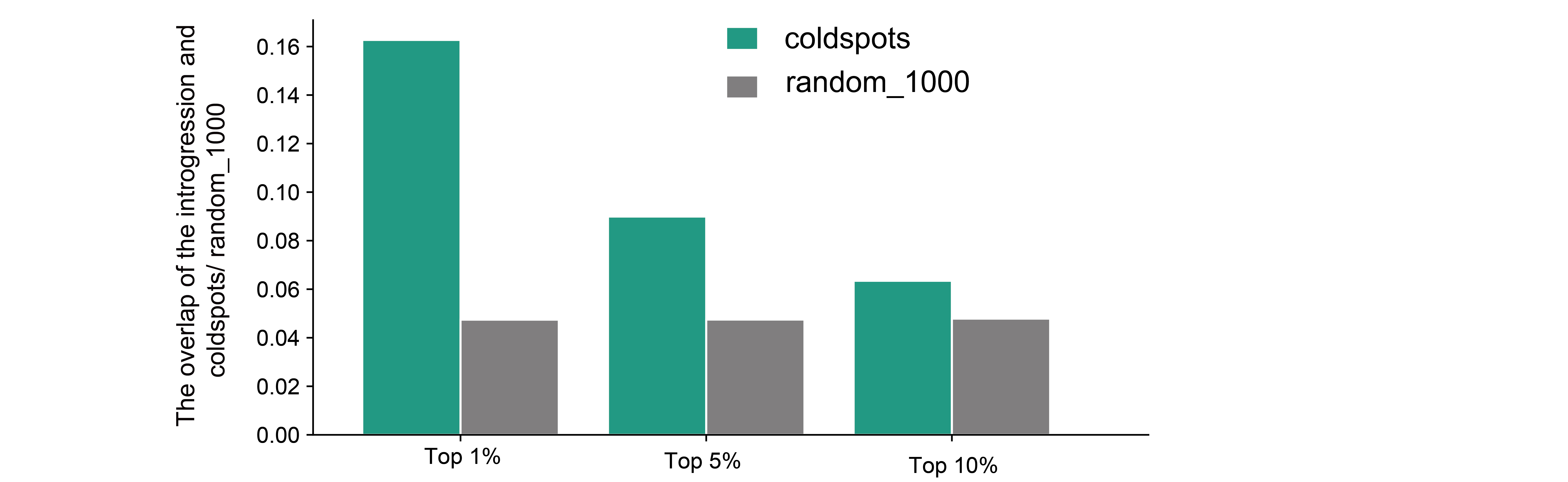

**Fig. S17.**  The overlap between recombination coldspots and introgressed regions in WL, as identified by Loter's top 1%, top 5%, and top 10% criteria, reveals a significantly higher concentration of coldspots within these introgressed regions compared to randomly selected genomic regions (n=1000).

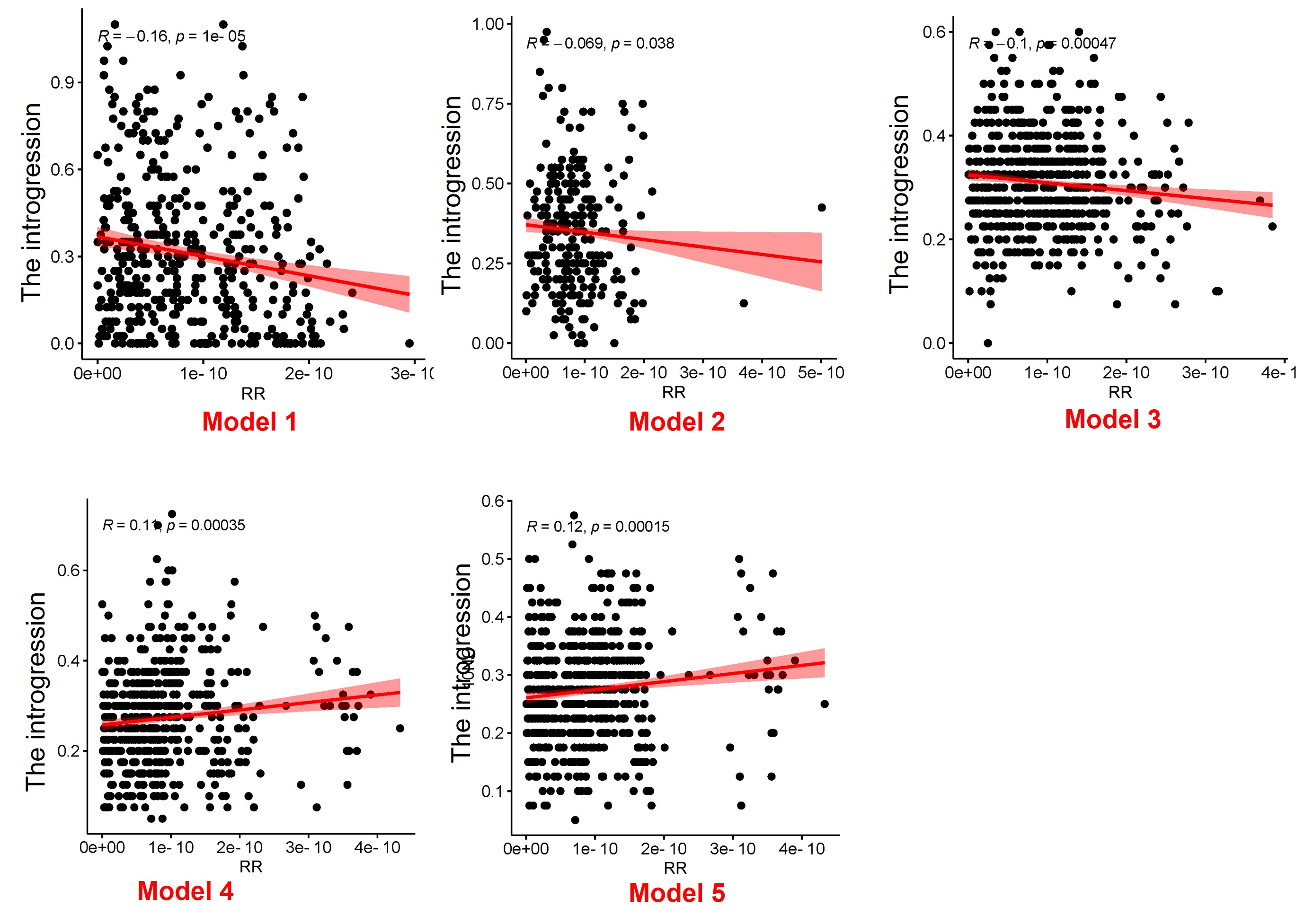

**Fig. S18.** The Pearson’s correlation coefficients between the frequency of introgression and recombination rate (c/bp) under five models.
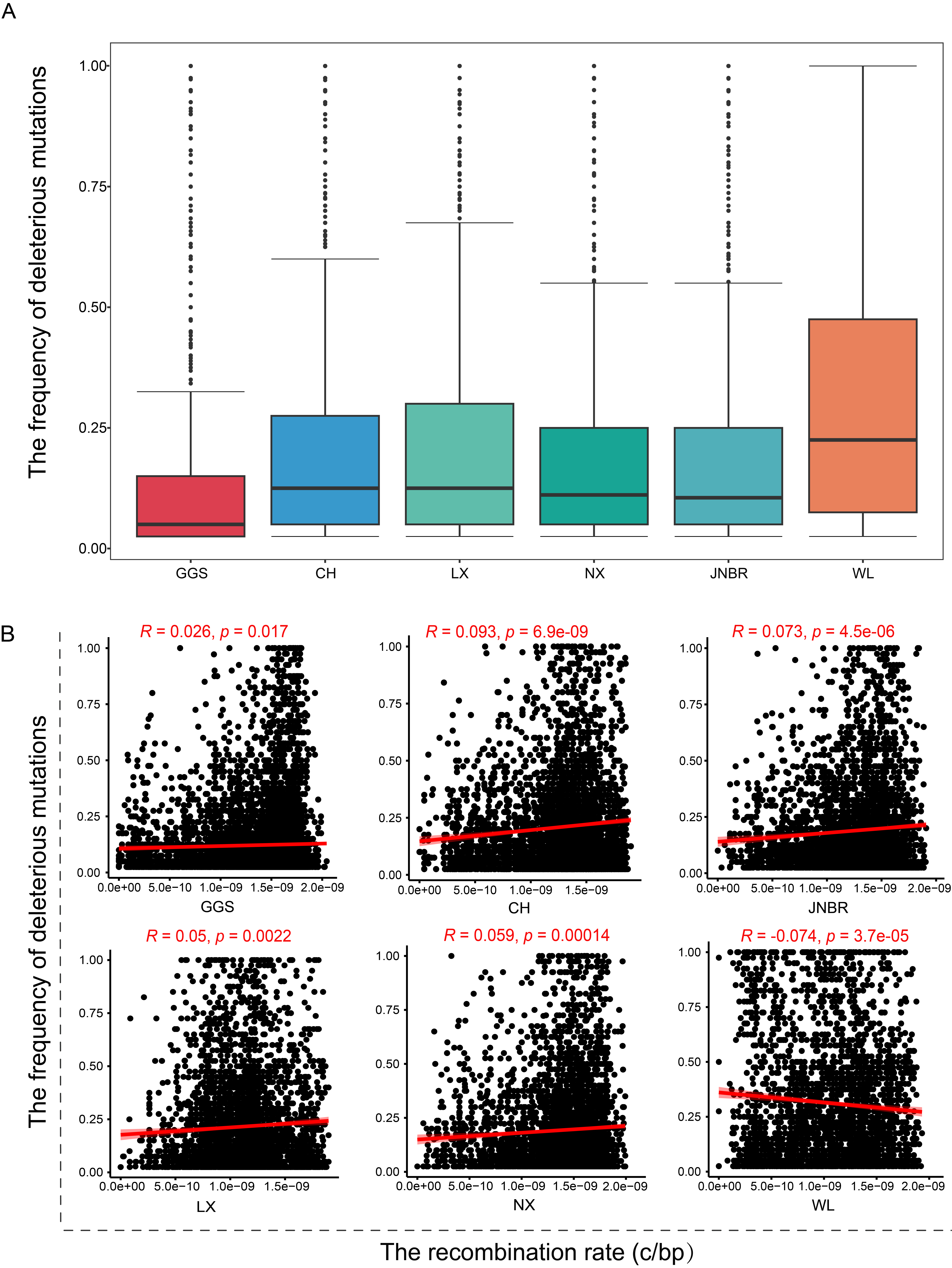

**Fig. S19.** (A) The frequency of deleterious mutations, and (B) the association between the recombination rate and the frequency of deleterious mutations.

**Supplementary Tables**

| **Table S1. Sample information of chicken** | | | | | | | | | |
| --- | --- | --- | --- | --- | --- | --- | --- | --- | --- |
| **Breed** | | **SampleName** | **SampleID** | | | | **Project** | | **Mean_Depth** |
| GGS | | GGSYP18774 | SAMC3562511 | | | | In this study | | 10.0339 |
| GGS | | GGSYP18878 | SAMC3562512 | | | | In this study | | 9.9703 |
| GGS | | GGSYP19900 | SAMC3562513 | | | | In this study | | 18.5946 |
| GGS | | GGSYP19908 | SAMC3562514 | | | | In this study | | 15.3439 |
| GGS | | GGSYPT110 | SAMC3562515 | | | | In this study | | 20.4573 |
| GGS | | GGSYPT114 | SAMC3562516 | | | | In this study | | 20.3495 |
| GGS | | GGSYPT119 | SAMC3562517 | | | | In this study | | 20.6312 |
| GGS | | GGSYPT130 | SAMC3562518 | | | | In this study | | 21.1463 |
| GGS | | GGSYPT137 | SAMC3562519 | | | | In this study | | 19.9988 |
| GGS | | GGSYPT147 | SAMC3562520 | | | | In this study | | 20.0261 |
| GGS | | GGSypt3011 | SAMC3562521 | | | | In this study | | 25.8346 |
| GGS | | GGSypt3016 | SAMC3562522 | | | | In this study | | 23.775 |
| GGS | | GGSypt3028 | SAMC3562523 | | | | In this study | | 23.3467 |
| GGS | | GGSypt3038 | SAMC3562524 | | | | In this study | | 24.7108 |
| GGS | | GGSypt3041 | SAMC3562525 | | | | In this study | | 26.1368 |
| GGS | | GGSypt3046 | SAMC3562526 | | | | In this study | | 24.6366 |
| GGS | | GGSypt3052 | SAMC3562527 | | | | In this study | | 24.7134 |
| GGS | | GGSypt3001 | ERR5036673 | | | | PRJEB42301 | | 23.6568 |
| GGS | | GGSypt3006 | ERR5036678 | | | | PRJEB42301 | | 25.1506 |
| GGS | | GGSypt3009 | ERR5036681 | | | | PRJEB42301 | | 27.6412 |
| JNBR | | JNBR.ypt5287 | SAMC3562528 | | | | In this study | | 9.67442 |
| JNBR | | JNBR.ypt5288 | SAMC3562529 | | | | In this study | | 10.3465 |
| JNBR | | JNBR.ypt5289 | SAMC3562530 | | | | In this study | | 9.99818 |
| JNBR | | JNBR.ypt5290 | SAMC3562531 | | | | In this study | | 9.76965 |
| JNBR | | JNBR.ypt5291 | SAMC3562532 | | | | In this study | | 8.49107 |
| JNBR | | JNBR.ypt5293 | SAMC3562533 | | | | In this study | | 10.1813 |
| JNBR | | JNBR.ypt5294 | SAMC3562534 | | | | In this study | | 9.48829 |
| JNBR | | JNBR.ypt5295 | SAMC3562535 | | | | In this study | | 10.6934 |
| JNBR | | JNBR.ypt5296 | SAMC3562536 | | | | In this study | | 8.9432 |
| JNBR | | JNBR.ypt5297 | SAMC3562537 | | | | In this study | | 9.17939 |
| JNBR | | JNBR.ypt5328 | SAMC3562538 | | | | In this study | | 7.89397 |
| JNBR | | JNBR.ypt5329 | SAMC3562539 | | | | In this study | | 9.35417 |
| JNBR | | JNBR.ypt5330 | SAMC3562540 | | | | In this study | | 8.20776 |
| JNBR | | JNBR.ypt5331 | SAMC3562541 | | | | In this study | | 9.53151 |
| JNBR | | JNBR.ypt5332 | SAMC3562542 | | | | In this study | | 9.14443 |
| JNBR | | JNBR.ypt5333 | SAMC3562543 | | | | In this study | | 9.61677 |
| JNBR | | JNBR.ypt5334 | SAMC3562544 | | | | In this study | | 8.85072 |
| JNBR | | JNBR.ypt5335 | SAMC3562545 | | | | In this study | | 9.69901 |
| JNBR | | JNBR.ypt5336 | SAMC3562546 | | | | In this study | | 9.37726 |
| JNBR | | JNBR.ypt5337 | SAMC3562547 | | | | In this study | | 8.78223 |
| WL | | WL10_FDSW202365605 | SAMC3562548 | | | | In this study | | 23.1722 |
| WL | | WL11_FDSW202365606 | SAMC3562549 | | | | In this study | | 19.5707 |
| WL | | WL12_FDSW202365607 | SAMC3562550 | | | | In this study | | 18.3166 |
| WL | | WL13_FDSW202365608 | SAMC3562551 | | | | In this study | | 19.4833 |
| WL | | WL14_FDSW202365609 | SAMC3562552 | | | | In this study | | 23.2465 |
| WL | | WL15_FDSW202365610 | SAMC3562553 | | | | In this study | | 20.2324 |
| WL | | WL16_FDSW202365611 | SAMC3562554 | | | | In this study | | 20.9783 |
| WL | | WL17_FDSW202365612 | SAMC3562555 | | | | In this study | | 20.0784 |
| WL | | WL18_FDSW202365613 | SAMC3562556 | | | | In this study | | 21.1995 |
| WL | | WL19_FDSW202365614 | SAMC3562557 | | | | In this study | | 19.5138 |
| WL | | WL1_FDSW202365596 | SAMC3562558 | | | | In this study | | 19.926 |
| WL | | WL20_FDSW202365615 | SAMC3562559 | | | | In this study | | 19.1657 |
| WL | | WL2_FDSW202365597 | SAMC3562560 | | | | In this study | | 19.823 |
| WL | | WL3_FDSW202365598 | SAMC3562561 | | | | In this study | | 19.2359 |
| WL | | WL4_FDSW202365599 | SAMC3562562 | | | | In this study | | 19.5338 |
| WL | | WL5_FDSW202365600 | SAMC3562563 | | | | In this study | | 21.573 |
| WL | | WL6_FDSW202365601 | SAMC3562564 | | | | In this study | | 19.191 |
| WL | | WL7_FDSW202365602 | SAMC3562565 | | | | In this study | | 19.008 |
| WL | | WL8_FDSW202365603 | SAMC3562566 | | | | In this study | | 19.539 |
| WL | | WL9_FDSW202365604 | SAMC3562567 | | | | PRJCA025697 | | 18.3231 |
| CH | | CH.gbe.SRR3145345 | SRR3145345 | | | | PRJNA309581 | | 6.69566 |
| CH | | CH01.SRR16992171 | SRR16992171 | | | | PRJNA782225 | | 12.3771 |
| CH | | CH02.SRR16992170 | SRR16992170 | | | | PRJNA782225 | | 11.999 |
| CH | | CH03.SRR16992169 | SRR16992169 | | | | PRJNA782225 | | 11.3434 |
| CH | | CH04.SRR16992168 | SRR16992168 | | | | PRJNA782225 | | 11.4692 |
| CH | | CH05.SRR16992167 | SRR16992167 | | | | PRJNA782225 | | 12.4018 |
| CH | | CH06.SRR16992166 | SRR16992166 | | | | PRJNA782225 | | 11.6885 |
| CH | | CH07.SRR16992165 | SRR16992165 | | | | PRJNA782225 | | 12.0471 |
| CH | | CH08.SRR16992164 | SRR16992164 | | | | PRJNA782225 | | 12.5674 |
| CH | | CH09.SRR16992163 | SRR16992163 | | | | PRJNA782225 | | 12.4842 |
| CH | | CH10.SRR16992161 | SRR16992161 | | | | PRJNA782225 | | 11.6183 |
| CH | | CH11.SRR16992091 | SRR16992091 | | | | PRJNA782225 | | 12.1719 |
| CH | | CH12.SRR10855145 | SRR10855145 | | | | PRJNA597842 | | 8.05644 |
| CH | | CH12.SRR16992090 | SRR16992090 | | | | PRJNA782225 | | 12.6911 |
| CH | | CH13.SRR16992089 | SRR16992089 | | | | PRJNA782225 | | 11.625 |
| CH | | CH14.SRR16992088 | SRR16992088 | | | | PRJNA782225 | | 11.8371 |
| CH | | CH15.SRR16992087 | SRR16992087 | | | | PRJNA782225 | | 11.7387 |
| CH | | CH16.SRR16992086 | SRR16992086 | | | | PRJNA782225 | | 11.4281 |
| CH | | CH17.SRR16992270 | SRR16992270 | | | | PRJNA782225 | | 11.298 |
| CH | | CH18.SRR16992269 | SRR16992269 | | | | PRJNA782225 | | 12.0458 |
| LX | | LX.game.CRR619078 | CRR619078 | | | | PRJCA010532 | | 13.2707 |
| LX | | LX.game.CRR619079 | CRR619079 | | | | PRJCA010532 | | 12.5949 |
| LX | | LX.game.CRR619080 | CRR619080 | | | | PRJCA01053 | | 11.3738 |
| LX | | LX.game.CRR619081 | CRR619081 | | | | PRJCA01053 | | 13.0141 |
| LX | | LX.game.CRR619082 | CRR619082 | | | | PRJCA01053 | | 13.2359 |
| LX | | LX.game.CRR619083 | CRR619083 | | | | PRJCA01053 | | 12.3717 |
| LX | | LX.game.CRR619085 | CRR619085 | | | | PRJCA01053 | | 11.0076 |
| LX | | LX.game.CRR619086 | CRR619086 | | | | PRJCA01053 | | 11.6343 |
| LX | | LX.game.CRR619088 | CRR619088 | | | | PRJCA01053 | | 11.2666 |
| LX | | LX.game.CRR619089 | CRR619089 | | | | PRJCA01053 | | 11.1011 |
| LX | | LX.game.CRR619090 | CRR619090 | | | | PRJCA010532 | | 10.1174 |
| LX | | LX.game.CRR619091 | CRR619091 | | | | PRJCA010532 | | 11.857 |
| LX | | LX.game.CRR619092 | CRR619092 | | | | PRJCA01053 | | 11.0147 |
| LX | | LX.game.CRR619094 | CRR619094 | | | | PRJCA01053 | | 11.2214 |
| LX | | LX.game.CRR619095 | CRR619095 | | | PRJCA010532 | | | 13.1898 |
| LX | | LX.game.CRR619096 | CRR619096 | | | PRJCA010532 | | | 11.3234 |
| LX | | LXFight.SAMC000221 | SAMC000221 | | | PRJCA000093 | | | 10.1824 |
| LX | | LXgame01.SRR12145471 | SRR12145471 | | | PRJNA627080 | | | 10.1294 |
| LX | | LXgame09.SRR12145462 | SRR12145462 | | | PRJNA627080 | | | 10.1414 |
| LX | | LXgame10.SRR12145461 | SRR12145461 | | | PRJNA627080 | | | 10.4437 |
| NX | | NX01.SRR16992260 | SRR16992260 | | | PRJNA782225 | | | 12.3957 |
| NX | | NX02.SRR16992249 | SRR16992249 | | | PRJNA782225 | | | 12.1355 |
| NX | | NX03.SRR16992238 | SRR16992238 | | | PRJNA782225 | | | 11.89 |
| NX | | NX04.SRR16992227 | SRR16992227 | | | PRJNA782225 | | | 11.9984 |
| NX | | NX05.SRR16992216 | SRR16992216 | | | PRJNA782225 | | | 11.9595 |
| NX | | NX06.SRR16992205 | SRR16992205 | | | PRJNA782225 | | | 12.3224 |
| NX | | NX07.SRR16992194 | SRR16992194 | | | PRJNA782225 | | | 11.6226 |
| NX | | NX08.SRR16992183 | SRR16992183 | | | PRJNA782225 | | | 11.7809 |
| NX | | NX09.SRR16992174 | SRR16992174 | | | PRJNA782225 | | | 12.3309 |
| NX | | NX10.SRR16992172 | SRR16992172 | | | PRJNA782225 | | | 12.2495 |
| NX | | NX11.SRR16992114 | SRR16992114 | | | PRJNA782225 | | | 13.2009 |
| NX | | NX12.SRR16992113 | SRR16992113 | | | PRJNA782225 | | | 12.2601 |
| NX | | NX13.SRR16992112 | SRR16992112 | | | PRJNA782225 | | | 12.5541 |
| NX | | NX14.SRR16992111 | SRR16992111 | | | PRJNA782225 | | | 12.779 |
| NX | | NX15.SRR16992110 | SRR16992110 | | | PRJNA782225 | | | 12.0711 |
| NX | | NX16.SRR16992109 | SRR16992109 | | | PRJNA782225 | | | 12.5851 |
| NX | | NX17.SRR16992108 | SRR16992108 | | | PRJNA782225 | | | 12.2468 |
| NX | | NX18.SRR16992106 | SRR16992106 | | | PRJNA782225 | | | 11.7046 |
| NX | | NX19.SRR16992105 | SRR16992105 | | | PRJNA782225 | | | 12.5133 |
| NX | | NX20.SRR16992104 | SRR16992104 | | | PRJNA782225 | | | 12.0943 |
| YOU | YOU01.SRR12145513 | | SRR12145513 | | | PRJNA627080 | | | 9.09839 |
| YOU | YOU02.SRR12145512 | | SRR12145512 | | | PRJNA627080 | | | 8.7605 |
| YOU | YOU03.SRR12145511 | | SRR12145511 | | | PRJNA627080 | | | 8.85915 |
| YOU | YOU04.SRR12145510 | | SRR12145510 | | | PRJNA627080 | | | 10.7695 |
| YOU | YOU05.SRR12145508 | | SRR12145508 | | | PRJNA627080 | | | 10.8034 |
| YOU | YOU06.SRR12145507 | | SRR12145507 | | | PRJNA627080 | | | 7.72524 |
| YOU | YOU07.SRR12145506 | | SRR12145506 | | | PRJNA627080 | | | 7.93467 |
| YOU | YOU08.SRR12145505 | | SRR12145505 | | | PRJNA627080 | | | 8.52072 |
| YOU | YOU09.SRR12145504 | | SRR12145504 | | | PRJNA627080 | | | 8.96798 |
| YOU | YOU10.SRR12145503 | | SRR12145503 | | | PRJNA627080 | | | 8.59662 |
| SK | SilKie.SRR12145425 | | SRR12145425 | | | PRJNA627080 | | | 8.9704 |
| SK | SilKie.SRR12145428 | | SRR12145428 | | | PRJNA627080 | | | 9.0842 |
| SK | SilKie.SRR12145429 | | | SRR12145429 | PRJNA627080 | | | 8.74547 | |
| SK | SilKie.SRR12145430 | | | SRR12145430 | PRJNA627080 | | | 8.52543 | |
| SK | SilKie.SRR12145432 | | SRR12145432 | | PRJNA627080 | | | 9.17776 | |
| SK | SilKie.SRR12145434 | | | SRR12145434 | PRJNA627080 | | | 8.53944 | |
| SK | SilKie01.SRR12145435 | | | SRR12145435 | PRJNA627080 | | | 9.58607 | |
| SK | SilKie04.SRR12145431 | | | SRR12145431 | PRJNA627080 | | | 8.12932 | |
| SK | SilKie08.SRR12145427 | | | SRR12145427 | PRJNA627080 | | | 8.25836 | |
| SK | SilKie11.SRR12145424 | | | SRR12145424 | PRJNA627080 | | | 8.10766 | |
| GV | Gallusvarius.SRR8362940 | | | SRR8362940 | PRJNA432200 | | | 29.9478 | |

|  | **Table S2. The SNP density of chicken** | | |
| --- | --- | --- | --- |
| **Breed** | | **The number of SNP** | **SNP densty** |
| WL | | 7,203,349 | 7.62287 |
| LX | | 11,422,726 | 12.0908 |
| CH | | 12,050,864 | 12.7522 |
| JNBR | | 12,197,010 | 12.9105 |
| NX | | 12,474,551 | 13.2004 |
| GGS | | 14,171,854 | 14.9954 |

| **Table S3. Sample information for calculating recombination rate in other species** | | | |
| --- | --- | --- | --- |
| **Sample ID** | **Project** | **Breed** | **Species** |
| SAMEA2612520 | PRJEB1683 | Asian wild boar | Pig |
| SAMEA2612521 | PRJEB1683 | Asian wild boar | Pig |
| SAMEA3497815 | PRJEB9922 | Asian wild boar | Pig |
| SAMEA3497816 | PRJEB9922 | Asian wild boar | Pig |
| SAMEA3497818 | PRJEB9922 | Asian wild boar | Pig |
| SAMEA3497819 | PRJEB9922 | Asian wild boar | Pig |
| SAMEA3497820 | PRJEB9922 | Asian wild boar | Pig |
| SAMEA3497821 | PRJEB9922 | Asian wild boar | Pig |
| SAMEA3497822 | PRJEB9922 | Asian wild boar | Pig |
| SAMEA3497823 | PRJEB9922 | Asian wild boar | Pig |
| SAMEA3497800 | PRJEB9922 | Meishan | Pig |
| SAMEA3497801 | PRJEB9922 | Meishan | Pig |
| SAMEA3497802 | PRJEB9922 | Meishan | Pig |
| SAMEA3497803 | PRJEB9922 | Meishan | Pig |
| SAMEA3497804 | PRJEB9922 | Meishan | Pig |
| SAMEA3497805 | PRJEB9922 | Meishan | Pig |
| SAMEA3497806 | PRJEB9922 | Meishan | Pig |
| SAMEA3497807 | PRJEB9922 | Meishan | Pig |
| SAMEA3497808 | PRJEB9922 | Meishan | Pig |
| SAMEA3497809 | PRJEB9922 | Meishan | Pig |
| SAMN02646525 | PRJNA238851 | Rongchang | Pig |
| SAMN02646543 | PRJNA238851 | Rongchang | Pig |
| SAMN02646545 | PRJNA238851 | Rongchang | Pig |
| SAMN04440482 | PRJNA309108 | Rongchang | Pig |
| SAMN05162341 | PRJNA322309 | Rongchang | Pig |
| SAMN06115544 | PRJNA322309 | Rongchang | Pig |
| SAMN06115552 | PRJNA322309 | Rongchang | Pig |
| SAMN06115553 | PRJNA322309 | Rongchang | Pig |
| SAMN06560025 | PRJNA378496 | Rongchang | Pig |
| SAMN06560051 | PRJNA378496 | Rongchang | Pig |
| SAMN11019701 | PRJNA524263 | Wannan Black | Pig |
| SAMN11019702 | PRJNA524263 | Wannan Black | Pig |
| SAMN11019703 | PRJNA524263 | Wannan Black | Pig |
| SAMN11019704 | PRJNA524263 | Wannan Black | Pig |
| SAMN11019705 | PRJNA524263 | Wannan Black | Pig |
| SAMN11019706 | PRJNA524263 | Wannan Black | Pig |
| SAMN11019707 | PRJNA524263 | Wannan Black | Pig |
| SAMN11019709 | PRJNA524263 | Wannan Black | Pig |
| SAMN11019710 | PRJNA524263 | Wannan Black | Pig |
| SAMN11019712 | PRJNA524263 | Wannan Black | Pig |
| ERR340330 | PRJEB3136 | Capra aegagrus from Iran | Goat |
| ERR340331 | PRJEB3136 | Capra aegagrus from Iran | Goat |
| ERR340334 | PRJEB3136 | Capra aegagrus from Iran | Goat |
| ERR340340 | PRJEB3136 | Capra aegagrus from Iran | Goat |
| ERR340345 | PRJEB3136 | Capra aegagrus from Iran | Goat |
| ERR340347 | PRJEB3136 | Capra aegagrus from Iran | Goat |
| ERR340426 | PRJEB3136 | Capra aegagrus from Iran | Goat |
| ERR470100 | PRJEB3136 | Capra aegagrus from Iran | Goat |
| ERR470104 | PRJEB3137 | Capra aegagrus from Iran | Goat |
| ERR470106 | PRJEB3138 | Capra aegagrus from Iran | Goat |
| ERR3281517 | PRJEB31857 | Boer | Goat |
| ERR3281518 | PRJEB31857 | Boer | Goat |
| ERR3281580 | PRJEB31857 | Boer | Goat |
| ERR3281581 | PRJEB31857 | Boer | Goat |
| ERR3281582 | PRJEB31857 | Boer | Goat |
| ERR3281583 | PRJEB31857 | Boer | Goat |
| ERR3284996 | PRJEB31857 | Boer | Goat |
| ERR3284997 | PRJEB31857 | Boer | Goat |
| ERR3284998 | PRJEB31857 | Boer | Goat |
| ERR3284999 | PRJEB31857 | Boer | Goat |
| SRR10083581 | PRJNA560446 | Jining_grey | Goat |
| SRR10083582 | PRJNA560446 | Jining_grey | Goat |
| SRR10083583 | PRJNA560446 | Jining_grey | Goat |
| SRR10083584 | PRJNA560446 | Jining_grey | Goat |
| SRR10083585 | PRJNA560446 | Jining_grey | Goat |
| SRR10083586 | PRJNA560446 | Jining_grey | Goat |
| SRR10083590 | PRJNA560446 | Jining_grey | Goat |
| SRR10083594 | PRJNA560446 | Jining_grey | Goat |
| SRR10083597 | PRJNA560446 | Jining_grey | Goat |
| SRR10083599 | PRJNA560446 | Jining_grey | Goat |
| SRR16990507 | PRJNA780399 | Shaanbei_white | Goat |
| SRR16990508 | PRJNA780399 | Shaanbei_white | Goat |
| SRR16990509 | PRJNA780399 | Shaanbei_white | Goat |
| SRR16990510 | PRJNA780399 | Shaanbei_white | Goat |
| SRR16990514 | PRJNA780399 | Shaanbei_white | Goat |
| SRR16990515 | PRJNA780399 | Shaanbei_white | Goat |
| SRR16990516 | PRJNA780399 | Shaanbei_white | Goat |
| SRR16990518 | PRJNA780399 | Shaanbei_white | Goat |
| SRR6040167 | PRJNA401972 | Mallard | Duck |
| SRR6323870 | PRJNA419832 | Mallard | Duck |
| SRR6323871 | PRJNA419832 | Mallard | Duck |
| SRR6323880 | PRJNA419832 | Mallard | Duck |
| SRR6323899 | PRJNA419832 | Mallard | Duck |
| SRR6323900 | PRJNA419832 | Mallard | Duck |
| SRR6323901 | PRJNA419832 | Mallard | Duck |
| SRR6323902 | PRJNA419832 | Mallard | Duck |
| SRR6323906 | PRJNA419832 | Mallard | Duck |
| SRR7091440 | PRJNA450892 | Mallard | Duck |
| SRR6323885 | PRJNA419832 | Gaoyou | Duck |
| SRR6323915 | PRJNA419832 | Gaoyou | Duck |
| SRR6323916 | PRJNA419832 | Gaoyou | Duck |
| SRR6323917 | PRJNA419832 | Gaoyou | Duck |
| SRR6323920 | PRJNA419832 | Gaoyou | Duck |
| SRR6323937 | PRJNA419832 | Gaoyou | Duck |
| SRR6323944 | PRJNA419832 | Gaoyou | Duck |
| SRR7091480 | PRJNA450892 | Gaoyou | Duck |
| SRR7091481 | PRJNA450892 | Gaoyou | Duck |
| SRR7091482 | PRJNA450892 | Gaoyou | Duck |
| SRR6323881 | PRJNA419832 | Jinding | Duck |
| SRR6323884 | PRJNA419832 | Jinding | Duck |
| SRR6323887 | PRJNA419832 | Jinding | Duck |
| SRR6323891 | PRJNA419832 | Jinding | Duck |
| SRR6323912 | PRJNA419832 | Jinding | Duck |
| SRR6323913 | PRJNA419832 | Jinding | Duck |
| SRR6323914 | PRJNA419832 | Jinding | Duck |
| SRR6323924 | PRJNA419832 | Jinding | Duck |
| SRR7091473 | PRJNA450892 | Jinding | Duck |
| SRR7091483 | PRJNA450892 | Jinding | Duck |
| SRR7091484 | PRJNA450892 | Jinding | Duck |
| SRR7091485 | PRJNA450892 | Jinding | Duck |
| SRR040294 | PRJNA46621 | Pekin | Duck |
| SRR6323904 | PRJNA419832 | Pekin | Duck |
| SRR7091411 | PRJNA450892 | Pekin | Duck |
| SRR7091412 | PRJNA450892 | Pekin | Duck |
| SRR7091417 | PRJNA450892 | Pekin | Duck |
| SRR7091419 | PRJNA450892 | Pekin | Duck |
| SRR7091432 | PRJNA450892 | Pekin | Duck |
| SRR7091488 | PRJNA450892 | Pekin | Duck |
| SRR7091500 | PRJNA450892 | Pekin | Duck |
| SRR7091501 | PRJNA450892 | Pekin | Duck |
| SRR7091505 | PRJNA450892 | Pekin | Duck |
| SRR7091506 | PRJNA450892 | Pekin | Duck |
| ERR157930 | PRJNA624020 | Moflon sheep | Sheep |
| ERR157931 | PRJNA624020 | Moflon sheep | Sheep |
| ERR157932 | PRJNA624020 | Moflon sheep | Sheep |
| ERR157935 | PRJNA624020 | Moflon sheep | Sheep |
| ERR157938 | PRJNA624020 | Moflon sheep | Sheep |
| ERR157939 | PRJNA624020 | Moflon sheep | Sheep |
| ERR157942 | PRJNA624020 | Moflon sheep | Sheep |
| ERR157944 | PRJNA624020 | Moflon sheep | Sheep |
| ERR466544 | PRJEB3139 | Moflon sheep | Sheep |
| ERR466545 | PRJEB3139 | Moflon sheep | Sheep |
| SRR16970290 | PRJNA781291 | Merino | Sheep |
| SRR24282371 | PRJNA781291 | Merino | Sheep |
| SRR24282372 | PRJNA781291 | Merino | Sheep |
| SRR24282373 | PRJNA781291 | Merino | Sheep |
| SRR24282374 | PRJNA781291 | Merino | Sheep |
| SRR24282375 | PRJNA781291 | Merino | Sheep |
| SRR24282376 | PRJNA781291 | Merino | Sheep |
| SRR24282377 | PRJNA781291 | Merino | Sheep |
| SRR24282378 | PRJNA781291 | Merino | Sheep |
| SRR24282379 | PRJNA781291 | Merino | Sheep |
| SRR11657475 | PRJNA624020 | Small-tail han | Sheep |
| SRR11657486 | PRJNA624020 | Small-tail han | Sheep |
| SRR11657497 | PRJNA624020 | Small-tail han | Sheep |
| SRR11657529 | PRJNA624020 | Small-tail han | Sheep |
| SRR11657572 | PRJNA624020 | Small-tail han | Sheep |
| SRR11657578 | PRJNA624020 | Small-tail han | Sheep |
| SRR11657584 | PRJNA624020 | Small-tail han | Sheep |
| SRR11657603 | PRJNA624020 | Small-tail han | Sheep |
| SRR11657681 | PRJNA624020 | Small-tail han | Sheep |
| SRR11657682 | PRJNA624020 | Small-tail han | Sheep |
| SAMN04306103 | PRJNA304478 | Tan sheep | Sheep |
| SAMN04306104 | PRJNA304478 | Tan sheep | Sheep |
| SAMN04306105 | PRJNA304478 | Tan sheep | Sheep |
| SAMN04306106 | PRJNA304478 | Tan sheep | Sheep |
| SAMN04306107 | PRJNA304478 | Tan sheep | Sheep |
| SAMN04306107 | PRJNA304478 | Tan sheep | Sheep |
| SAMN04306108 | PRJNA304478 | Tan sheep | Sheep |
| SAMN04306112 | PRJNA304478 | Tan sheep | Sheep |
| SAMN04306112 | PRJNA304478 | Tan sheep | Sheep |
| SAMN04306112 | PRJNA304478 | Tan sheep | Sheep |
